# Transposable Elements Shape Stemness in Normal and Leukemic Hematopoiesis

**DOI:** 10.1101/2021.02.16.431334

**Authors:** Giacomo Grillo, Bettina Nadorp, Aditi Qamra, Amanda Mitchell, Christopher Arlidge, Ankita Nand, Naoya Takayama, Alex Murison, Seyed Ali Madani Tonekaboni, Andrea Arruda, Jean C. Y. Wang, Mark D. Minden, Özgen Deniz, Héléna Boutzen, John E. Dick, Mathieu Lupien

**Author notes:** Equal contribution (Giacomo Grillo, Bettina Nadorp). Corresponding author: John E. Dick and Mathieu Lupien.

## Abstract

Despite most acute myeloid leukemia (AML) patients achieving complete remission after induction chemotherapy, two-thirds will relapse with fatal disease within five years. AML is organized as a cellular hierarchy sustained by leukemia stem cells (LSC) at the apex, with LSC properties directly linked to tumor progression, therapy failure, and disease relapse ^1–5^. Despite the central role of LSC in poor patient outcomes, little is known about the genetic determinants driving their stemness properties. As LSCs share many functional and molecular properties with normal hematopoietic stem cells (HSC) ^6^, we investigated accessible chromatin unique across normal hematopoietic and cancer cell states and identified transposable elements (TEs) as genetic determinants of both primitive populations in comparison with their downstream mature progeny. A clinically-relevant TE chromatin accessibility-based LSCTE121 signature was developed that enabled patient classification based on survival outcomes. Through functional assays, primitive cell specific-TE subfamilies were found to serve as docking sites for stem cell-associated regulators of genome topology or lineage-specific transcription factors, including LYL1 in LSCs. Finally, using chromatin editing tools, we establish that chromatin accessibility at LTR12C elements in LSCs are necessary to maintain stemness properties. Our work identifies TEs as genetic drivers of primitive versus mature cell states, where distinct TE subfamilies account for stemness properties in normal versus leukemic hematopoietic stem cells.

## Main

Lifelong blood production originates from self-renewing, multipotent hematopoietic stem cells (HSCs) that give rise to the full hierarchy of progenitor and then mature hematopoietic populations ^7^. The phenotypic heterogeneity across the hematopoietic hierarchy relies on population-specific genetic architectures, where distinct populations arise from the contribution of different sections of a shared genome ^8–11^. For example, comparing chromatin accessibility across hematopoietic populations identified chromatin variants, defined as genetic elements varying in chromatin accessibility between populations along the differentiation axis ^12^, and revealed a role for CTCF in mediating changes to the three-dimensional genome organization to govern the transition of HSC from dormant, low primed to activated, primed states ^13^. By contrast, chromatin variants specific to individual mature populations serve as cis-regulatory elements by acting as binding sites for lineage-specific transcription factors and promoting lineage-specific expression profiles ^13,14^. Hence, identifying chromatin variants across hematopoietic populations reveals the DNA sequences undergoing chromatin reprogramming to drive phenotypic heterogeneity.

Akin to normal hematopoiesis, acute myeloid leukemia (AML) is hierarchically organized ^6^, with leukemia stem cells (LSCs) at the apex ^1,2^. Because LSCs possess self-renewal capacity, they sustain the long-term generation of malignant populations and represent the source of evolutionary diversity, driving leukemic progression, therapy failure and relapse ^3–5^. The clinical relevance of LSCs has been established in two ways. First, a LSC-specific gene expression signature score (LSC17) was developed and found to accurately predict both overall patient survival and response to standard induction therapy ^15,16^. Second, mutational tracing studies in paired diagnosis/relapse AML samples showed the existence of genetically diverse, LSC-driven subclones present in AML diagnosis samples, where some LSC are already fated to survive induction therapy and fuel relapse ^4^. Finally, the link between HSCs and LSCs during human leukemogenesis has also been made. Mutations that give rise to clonal hematopoiesis are amongst the first changes to the genetic architecture of HSCs that create a pre-leukemic HSC reservoir that further evolves leading to LSCs and fulminant AML ^4,17,18^. However, we know little of the identity of additional non-genetic events such as chromatin variants that drive the transformation of HSCs into LSCs. Although there are examples where non-genetic events have been broadly linked to cancer stem cell properties ^3,19,20^, little is known of the specific features of such determinants. Thus, a clear understanding of the non-genetic events found in LSCs that are distinct from HSCs is required to better understand how cancer stemness is governed and to devise antagonistic strategies to improve therapies by ensuring that LSC are eradicated.

To date, the search for chromatin variants that represent determinants of phenotypic heterogeneity across normal hematopoietic populations and AML has mainly focused on the non-repetitive genome, but this excludes transposable elements (TEs), which compose ∼50% of the human genome ^21,22^. However, TEs account for diverse genetic functions relevant to development and pathogenesis ^23^. TEs are classified in subfamilies composed of tens to thousands of individual elements traced back to a single ancestral unit ^24^. In AML, as in many other cancer types, the expression of specific subfamilies is repressed to prevent activation of the viral mimicry response otherwise activated by the double-stranded RNAs they encode ^25–27^. Other TE subfamilies lie in cancer-associated chromatin variants, where they function as *cis*-regulatory elements rather than encoding transcripts ^28,29^. Such TE subfamilies potentiate the expression of neighboring genes and/or impact the three-dimensional genome organization by recruiting sequence-specific transcription factors, including CTCF ^30–32^. This results in TE mediated oncogene expression in AML, as in various other cancer states ^29,31,33^. Despite their reported contribution to cancer, the role of TEs as determinants of stemness properties, such as in LSCs, remains unknown. In the present study, we show how different TE subfamilies are enriched in chromatin variants identified across the continuum of both normal hematopoietic and LSC-driven leukemic hierarchies to modulate stemness properties.

## Results

### Chromatin variants from primitive and mature populations contain distinct TE subfamilies

To systematically study TE subfamily enrichment within accessible chromatin across hematopoietic populations, we used the Assay for Transposase-Accessible Chromatin Sequencing (ATAC-Seq; see Methods) data generated across six highly purified hematopoietic stem and progenitor populations (HSPC), including long-term (LT) and short-term (ST) HSCs, common myeloid progenitors (CMP), megakaryocyte-erythroid progenitors (MEP), multi-lymphoid progenitors (MLP) and granulocyte-monocyte progenitors (GMP), as well as seven mature cell populations exhibiting myeloid and lymphoid fates isolated from human cord blood donors (Extended Data Fig. 1A) ^13^. These populations were selected according to the sorting scheme we developed (see Methods), which achieves the highest purity compared to other studies ^14^ and allows the isolation of HSCs with distinct repopulation and self-renewal potential (∼30%) for improved resolution at the apex of the normal hematopoietic hierarchy (see Methods). We selected chromatin accessibility because it enables the identification of chromatin variants that define cell state identity ^12–14,34,35^.

The enrichment of all TE subfamilies (n=971, hg38, see Methods) within the accessible chromatin landscape of hematopoiesis was calculated using ChromVAR (see Methods). We used the genomic coordinates of TE subfamily elements instead of transcription factor DNA recognition sequence positions (see details in Methods) while still controlling for GC content. Unsupervised clustering of accessibility Z-scores for all TE subfamilies in different hematopoietic populations revealed that LT-, ST-HSCs and later progenitor populations, collectively referred to as HSPC, share similar accessible TE subfamily enrichment profiles and collectively, these are distinct from mature populations (Fig. 1A). The megakaryocyte-erythroid lineage is the exception, with MEPs clustering with erythroid precursors that are more similar to other mature rather than HSPC populations (Fig. 1A) and could be caused by a strong lineage-specific enrichment of TE subfamilies. To show that the megakaryocyte-erythroid lineage-specific TE subfamilies are driving the separation in the unsupervised clustering, we identified the strongest megakaryocyte-erythroid-specific TE subfamilies (Benjamini-Hochberg corrected p-value (q-value)<0.0001, Extended Data Fig. 1B) and subsequently removed them from the list of TE subfamilies used in unsupervised clustering. This resulted in a clear-cut separation of HPSC versus mature populations (Extended Data Fig. 1C). Thus, our data demonstrate that the chromatin variants reported from chromatin accessibility data across specific TE subfamilies distinguish HSPC from mature hematopoietic cells.

**Fig. 1:**
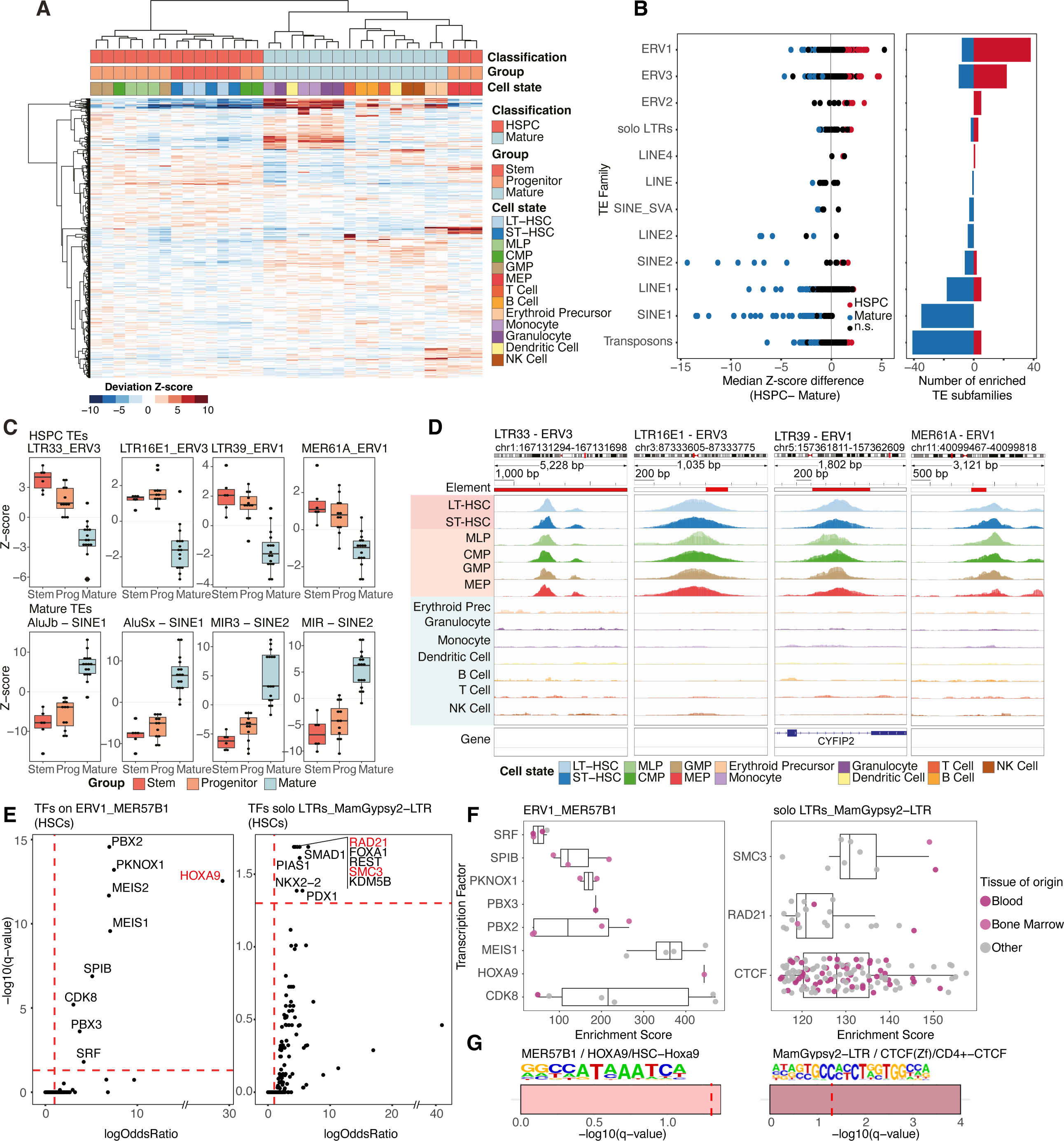
Distinct TE subfamilies populate the accessible chromatin of primitive and mature hematopoietic cells. A) Heatmap displaying differential accessibility Z-scores of all TE subfamilies across bulk ATAC-seq profiles of HSC, progenitor and mature hematopoietic populations; note that HSC and progenitors show similar TE subfamily enrichment profiles. B) Association of TE families with ’stemness’. Direct comparison of HSC and progenitor (“primitive”) versus mature cells. Y-axis shows all TE families with enriched TE subfamilies (each dot denotes a different TE subfamily). LINE1, SINE1, SINE2, and Transposons subfamilies are enriched in mature, whereas ERV1 and ERV3 families are enriched in HSPC cells (left). Number of enriched TE subfamilies in HSPC or mature cells, divided by TE family (right). TE families are ordered from most highly enriched in HSPC to mature cells. n.s., not significant. C) Examples of HSPC and mature enriched TE subfamilies of the ERV1, ERV3, SINE1 and SINE2 TE families. Boxplots show differential accessibility Z-scores in hematopoietic stem, progenitor and mature populations. D) Genome browser view of ATAC-seq signal from hematopoietic populations for one element of each primitive enriched TE subfamily shown in C). E) Transcription factor cistromes enrichment at 2 of the 23 TE subfamilies enriched in stem populations (HSCs). Every dot corresponds to a transcription factor cistrome. Red dashed lines correspond to -log10(q)=1.3 (q=0.05) and logOddsRatio(logOR)=1 thresholds. F) Individual HOXA9 (left) or three-dimensional genome organization factor (right) cistromes enriched over HSC-specific TE subfamilies (according to Fig. 1E). Box plots showing enrichment GIGGLE scores of individual cistromes profiled in cell lines derived from a variety of tissue states. G) Enrichment of HOXA9 and CTCF DNA recognition sequences within TE subfamilies enriched in HSCs. Bars represent -log10(q) for each TE subfamily for transcription factor DNA recognition sequences. The red dashed line corresponds to -log10(q)=1.3 (q=0.05) threshold. Abbreviations: Long-term/Short-Term hematopoietic stem cells (LT-/ST-HSC), common myeloid progenitors (CMP), megakaryocyte-erythroid progenitors (MEP), myeloid-lymphoid progenitors (MLP), granulocyte-monocyte progenitor (GMP).

To identify which differentially accessible TE subfamilies are distinct between HSPC versus mature populations, we compared the accessibility Z-scores of all TE subfamilies using a two-sided Wilcoxon signed rank test. We found 81 and 128 TE subfamilies belonging to 12 TE families were enriched in the HSPC and mature populations, respectively (q < 0.01, Fig. 1B, Extended Data Fig. 1D, Extended Data Table 1). HSPC populations were mainly enriched in subfamilies belonging to ERV1 and ERV3 TE families (Fig. 1B). Enriched ERV1 subfamilies included MER61A (q=1.10e-05) and LTR39 (q=1.87e-06), while enriched ERV3 subfamilies included LTR16E1 (q=4.61e-07) and LTR33 (q=9.05e-07) (Fig 1C, Extended Data Fig. 1E), exemplified at unique elements (Fig. 1D). In contrast, mature populations were mainly enriched in SINE1, SINE2 and Transposon subfamily members (Fig. 1B), such as AluJB (q=1.26e-07) and AluSx (q=1.08e-07) of the SINE1 family, as well as MIR (q=2.93e-07) and MIR3 (q=2.93e-07) of the SINE2 family (Fig. 1C, Extended Data Fig. 1E). Despite finding a similar number of accessible DNA elements (aka: peaks) in each mature population (Extended Data Fig. 1F), the enrichment of SINE1, SINE2 and Transposons subfamily members relied on population-specific accessible elements. This contrasts with HSPC, where the same elements from ERV1, ERV3 subfamily members are accessible across all stem and progenitor populations. In conclusion, our study demonstrates distinct chromatin accessibility patterns of TE subfamilies in HSPC versus mature populations, highlighting their potential role in cell state identity within hematopoiesis.

### Accessible TE subfamilies that serve as docking sites for stem factors define primitive hematopoietic populations

We next assessed if chromatin variants over TE subfamilies could distinguish LT-HSCs (the most primitive HSC subset) from any other hematopoietic population (Extended Data Fig. 1A). Comparison of the accessibility Z-scores of all TE subfamilies did not identify differentially-enriched TE subfamilies across all one-to-one comparisons (Extended Data Table 2). We next compared stem versus progenitors and mature populations. Comparison of stem populations against the collection of mature populations revealed 61 TE subfamilies enriched in stem populations versus 119 in mature populations (q < 0.01, Extended Data Fig. 2A, Extended Data Table 3), while comparison of progenitor to mature populations identified 56 TE subfamilies enriched in progenitor versus 89 in mature populations (q < 0.01, Extended Data Table 4). Of all accessible TE subfamilies found to be distinct between HSPC compared to mature populations (Fig. 1A), twenty-three TE subfamilies were specific to the stem populations (Extended Data Fig. 2B-C). Interestingly, LTR85a, MLT1E2, and HERV4_I-int were the only two also enriched in stem populations compared to progenitor populations (q < 0.05, Extended Data Fig. 2D, Extended Data Table 5). Notably, MER4B-int, MER57B1, MER57B2, MER57-int, LTR16A2, LTR67B, and LTR85a TE subfamilies showed the highest Z-scores in the LT/HSPC and/or Act/HSPC signatures of chromatin accessibility that we previously showed could discriminate between LT-HSC, ST-HSC and other progenitor populations ^13^ (Extended Data Table 6). Collectively, our results reveal that chromatin variants over specific TE subfamilies discriminate HSPC from mature populations, with a limited set of twenty-three that are unique to the stem populations.

Each TE subfamily harbors DNA recognition sequences allowing them to recruit transcription factors and guide cell differentiation ^29,30,36–39^. We used the ReMap atlas of 485 transcription factor cistromes (see Methods) to examine the enrichment of these sequences over the twenty-three TE subfamilies that were specific to stem populations. We found enrichment of HOXA9, a well-known HSC factor ^40^, at elements of the MER57B1 (q=2.88e-13, logOR=27.69) and MER57B2 (q=0.0007, logOR=11.95) (Fig. 1E, Extended Data Fig. 2E) subfamilies. The cistrome for RUNX1, another well-known HSCs transcription factor ^41^, was found to be enriched over elements of the MLT1E2 (q=0.03, logOR=2.12) (Extended Data Fig. 2F). While none of the 485 cistromes in ReMap were enriched over HERV4_I-int, HERVK9-int, LTR80A, LTR85a, LTR9A1, MamGypLTR1b, MER127, MER4B-int, MER57-int, MER83B-int, MLT2E and PRIMA41-int elements, the cistromes of CTCF and/or cohesin complex factors (SMC1A2, SMC3 and RAD21), known regulators of genome topology ^42^, were enriched over MamGypsy2-LTR and LTR16A2 elements (Fig. 1E, Extended Data Fig. 2G, Extended Data Table 7). We validated the enrichment of these stem transcription factors by computing an independent score, the GIGGLE enrichment score ^43^. Our analysis revealed enrichment of HOXA9 cistrome over MER57B1 and MER57B2 elements (Fig. 1F, Extended Data Fig. 2H), enrichment of RUNX1 cistrome over MLT1E2 elements (Extended Data Fig. 2I) and enrichment of cistromes from CTCF and cohesin complex factors over MamGypsy2-LTR and LTR16A2 elements (Fig. 1F, Extended Data Fig. 2J). In parallel, we performed sequence analysis and identified the enrichment of the HOXA9 or CTCF DNA binding motifs within MER57B1 or MamGypsy2-LTR elements, respectively (Fig. 1G). Collectively, these results suggest that TE subfamilies uniquely accessible in stem populations provide docking sites for known “stem” transcription factors, including HOXA9, RUNX1, and components of the three-dimensional genome organization (CTCF, cohesin).

### LSCs exhibit HSPC-like TE subfamily enrichment within accessible chromatin

Considering the role of LSCs in AML etiology, we next examined whether chromatin variants over TE subfamilies could be used to discriminate populations enriched for self-renewing LSCs from leukemic cell fractions lacking LSCs as well as from normal HSPC and mature hematopoietic populations. Uncultured AML patient samples (n=15) were sorted into fractions based on CD34 and CD38 expression, and fractions were functionally classified as LSC+ (n=11) or LSC-(n=24) based on their ability to generate a leukemic graft upon xenotransplantation into immunodeficient mice, as described previously ^15,44^. In parallel, we performed ATAC-Seq and measured TE subfamily enrichment within the accessible chromatin profiles of LSC- and LSC+ fractions. Unsupervised clustering of Z-scores of all TE subfamilies in LSC+ and LSC-fractions, as well as normal hematopoietic populations, showed clustering of all LSC+ fractions with the HSPC populations (Fig. 2A). In contrast, most LSC-fractions (17/24) clustered not with LSC+ fractions but rather with mature hematopoietic populations (Fig. 2A). The discordance between the TE subfamily enrichment profile classification (as HSPC-like) and the functional classification (as LSC-) for the 7 LSC-fractions that clustered with the LSC+ fractions (2 CD34+/CD38-, 4 CD34+/CD38+, 1 CD34-/CD38+; Fig. 2A-B) may draw from low LSC frequency within these populations. In conclusion, our analysis reveals an association between chromatin variants over TE subfamilies and the discrimination of LSC populations, providing a nuanced classification for AML biology.

**Fig. 2:**
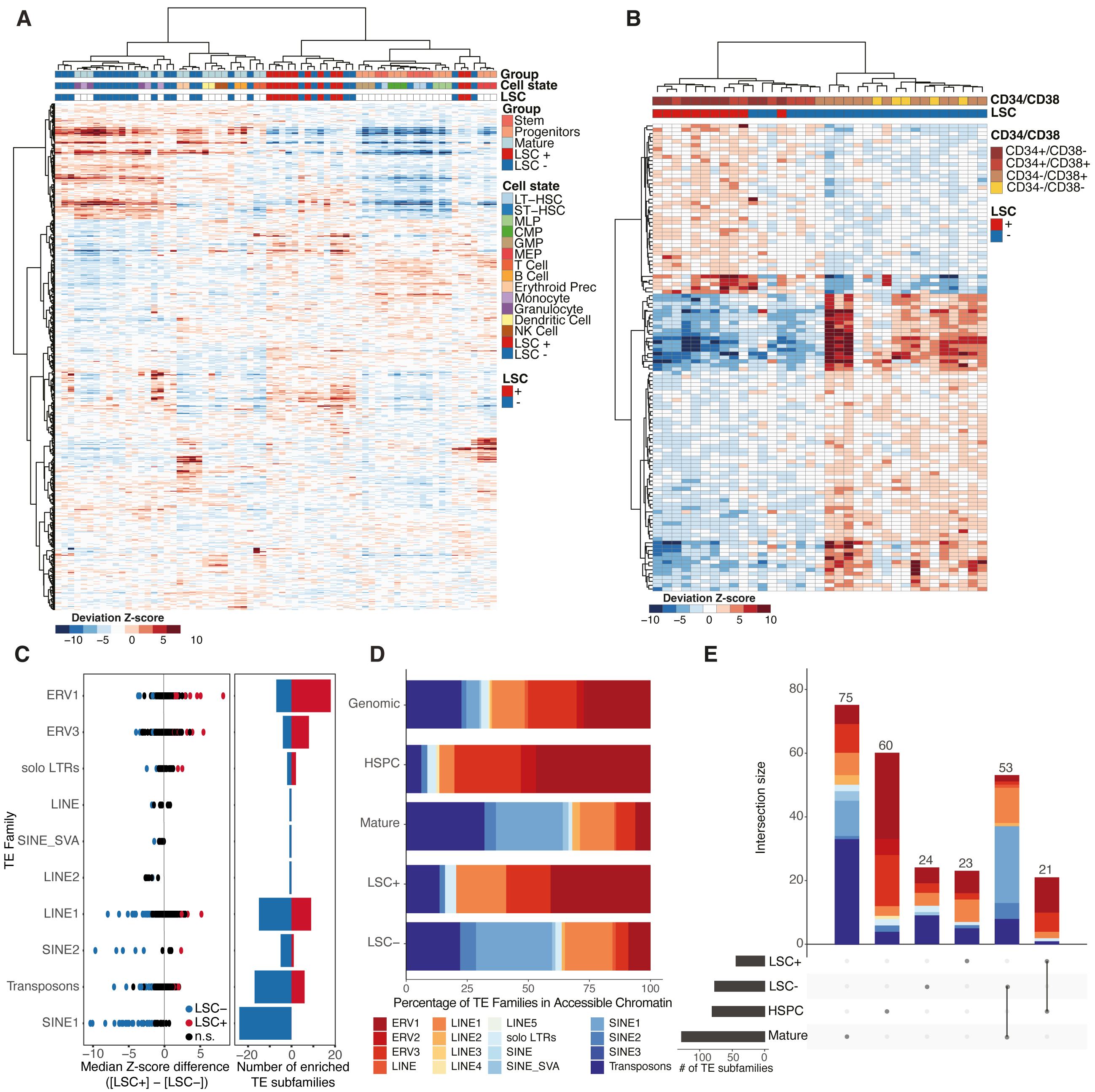
Leukemia stem cells containing fractions cluster with HSPCs and show distinct and common subfamily enrichment. A) Heatmap of accessibility Z-scores of all TE subfamilies (rows) across bulk ATAC-seq profiles of HSPC, mature hematopoietic populations, LSC+, and LSC-fractions clustered based on correlation. LSC+ fractions clustered with HSPC while LSC-fractions mostly clustered with mature populations. B) Heatmap of TE subfamilies with differential accessibility Z-scores (rows, q < 0.01) between ATAC-seq profiles of LSC+ (n=11) and LSC-fractions (n=23). C) Enrichment of TE families across LSC+ versus LSC-samples. The y-axis shows all TE families; points denote TE subfamilies. SINE1, SINE2, and Transposons families are enriched in LSC-samples, whereas ERV1 and ERV3 families are enriched in LSC+ samples. Number of enriched TE subfamilies in LSC+ or LSC-samples, divided by TE family (right). TE families are ordered from most highly enriched in LSC+ to LSC-. n.s., not significant. D) Comparison of the proportion of TE families enriched in HSPC and hematopoietic mature populations in LSC- and LSC+ fractions and the overall distribution of TE families within the hg38 build of the human genome. E) UpSet plot showing the intersection of TE subfamilies enriched in LSC+/HSPC (21 families) and LSC-/mature hematopoietic populations (53 families). Colors within bar graphs denote TE families as annotated in panel D.

Direct comparison of accessibility Z-scores of all TE subfamilies between LSC+ and LSC-fractions revealed 44 significantly enriched accessible TE subfamilies in LSC+ and 77 in LSC-fractions (Fig. 2B-C, Extended Data Fig. 3A-B, Extended Data Table 8). LSC+ fractions were mostly enriched for ERV1 and ERV3 families, whereas LSC-fractions showed enrichment of SINE1, SINE2, and Transposon families (Fig. 2C). Next, we compared accessible TE families and subfamilies across normal hematopoietic populations and LSC fractions. Overall, the proportions of TE families discriminating HSPC vs mature and LSC+ vs LSC-fractions were skewed compared to the proportions of TEs distributed across the human genome (Fig. 2D). Specifically, accessible TE families enriched in HSPC populations and LSC+ fractions consisted preferentially of ERV1 and ERV3 families (Fig. 2D). In contrast, accessible TE families enriched in mature populations and LSC-fractions had higher representation of SINE1 and Transposons families (Fig. 2D). The majority of TE subfamilies enriched in the accessible chromatin of LSC+ or LSC-fractions were shared with HSPC (21/44) or mature populations (53/77), respectively (Fig. 2E). In line with the similarities observed in the overall TE family distribution, the majority of common subfamilies between LSC+ fractions and HSPC populations (14/21 subfamilies) belonged to the ERV1 and ERV3 families (Fig. 2E, Extended Data Fig. 3C-D). In contrast, the majority of common subfamilies between LSC-fractions and mature populations belonged to the SINE1 and Transposons families (Fig. 2E, Extended Data Fig. 3C-D). Z-score comparisons of the enrichment in the common subfamilies between HSPC/LSC+ or mature/LSC-showed largely concordant enrichment (Extended Data Fig. 3D). Two TE subfamilies enriched in LSC+, but not HSPC accessible chromatin, namely LTR12C (ERV1 family) and L1PA7 (LINE1 family), showed the largest discordance between HSPC and LSC+ populations (Extended Data Fig. 3E). Although not discriminating LSC+ from LSC-fractions, all LINE1 families enriched in HSPC versus mature hematopoietic populations were more enriched in LSC+ compared to HSPC populations (Extended Data Fig. 3E); in line with the DNA hypomethylation over LINE1 elements reported across many cancers ^45^. Collectively, our data suggest that LSCs can be discriminated from committed leukemic cells based on the enrichment of specific TE subfamilies within their accessible chromatin that are largely shared with normal HSPC populations.

### An accessible TE subfamily-based signature shows clinical value as a marker of relapse in AML

Stemness is related to survival and relapse in AML and other cancer types ^46,47^. We hypothesized that TE subfamilies enriched in LSC+ and LSC-fractions (44 and 77 families enriched in LSC+ and LSC-respectively, as shown in Fig 2B-C, hereafter referred to as LSCTE121) could serve as a signature for stemness in AML and accordingly that this would be associated with clinical outcomes. To this end, we performed ATAC-seq on peripheral blood samples from two independent cohorts of AML patients (cohort1, n=29; cohort2, n=60) (Extended Data Table 9) and assessed each sample for their LSCTE121 signature score. Unsupervised clustering based on the LSCTE121 score grouped 9 of the cohort1 AML samples with the LSC-fractions whereas the other 20 AML samples clustered in the same main branch as the LSC+ fractions (Extended Data Fig. 4A). Similarly, 20 of the cohort2 AML samples clustered with the LSC-fractions whereas the other 40 clustered in the same main branch as the LSC+ fractions (Extended Data Fig. 4B). This classification of bulk AML samples into LSC-like signature score and non-LSC-like signature score is reminiscent of recent gene expression-based patient stratification, lending credence to our approach ^48,49^. To simplify the interpretation of TE subfamily enrichment, we created a combined TE Z-score restricted to the LSCTE121. This score uses Stouffer’s method to combine the enrichment of LSC+ and the depletion of LSC-TE subfamilies (see Methods). Comparison of the combined LSCTE121 Z-scores of samples in the major branches of the heatmaps (Extended Data Fig. 4A-B) showed the highest scores in the branch containing the LSC+ fractions (red branch). As expected, LSC+ fractions had higher LSCTE121 Z-scores than LSC-fractions (Extended Data Fig. 4C). Further, there was no correlation between the blast count of a given sample and the LSCTE121 Z-score in both cohorts (cohort1: R=0.031, p=0.88; cohort2: R=-0.17, p=0.21; Extended Data Fig. 4D-E), suggesting that the combined LSCTE121 Z-score stratification is a reflection of stemness and not of blast count. Kaplan-Meier estimates comparing the top 25th percentile with the bottom 25th percentile of LSCTE121 Z-score AML patients from cohort1 showed significant differences in disease-free and overall survival (Fig. 3A-C, Extended Data Fig. 4F), with in high and low LSCTE121 subgroups. Additionally, when compared with the LSC17 score, a gene expression-based score shown to predict overall patient survival and response to standard induction therapy ^15^, there was variation in how these two scoring methods classified samples ^15^ (Fig. 3A and 3D). We validated the prediction of overall survival in our patient cohort by using the LSC17 score (Extended Data Fig. 4G, log-rank p-value=2e-04) and we then sought to determine the similarity between the LSCTE121 Z-score and the LSC17 score. We observed low correlation between the LSC17 score and the LSCTE121 Z-score (R=0.27, p=0.16; Fig. 3D). However, the LSCTE121 score was significantly higher in fractions with the highest LSC frequency, assessed with limiting dilution assays, supporting the capture of stemness properties by the LSCTE121 Z-score (Fig. 3E). Furthermore, significant differences in disease-free survival were also observed with cohort2 AML patients (Fig. 3F-H, Extended Data Fig. 4H). Collectively, our results suggest that the LSCTE121 Z-score captures a different underlying biology than the LSC17 score and that each score provides risk stratification based on complementary discriminators of stemness.

**Fig. 3:**
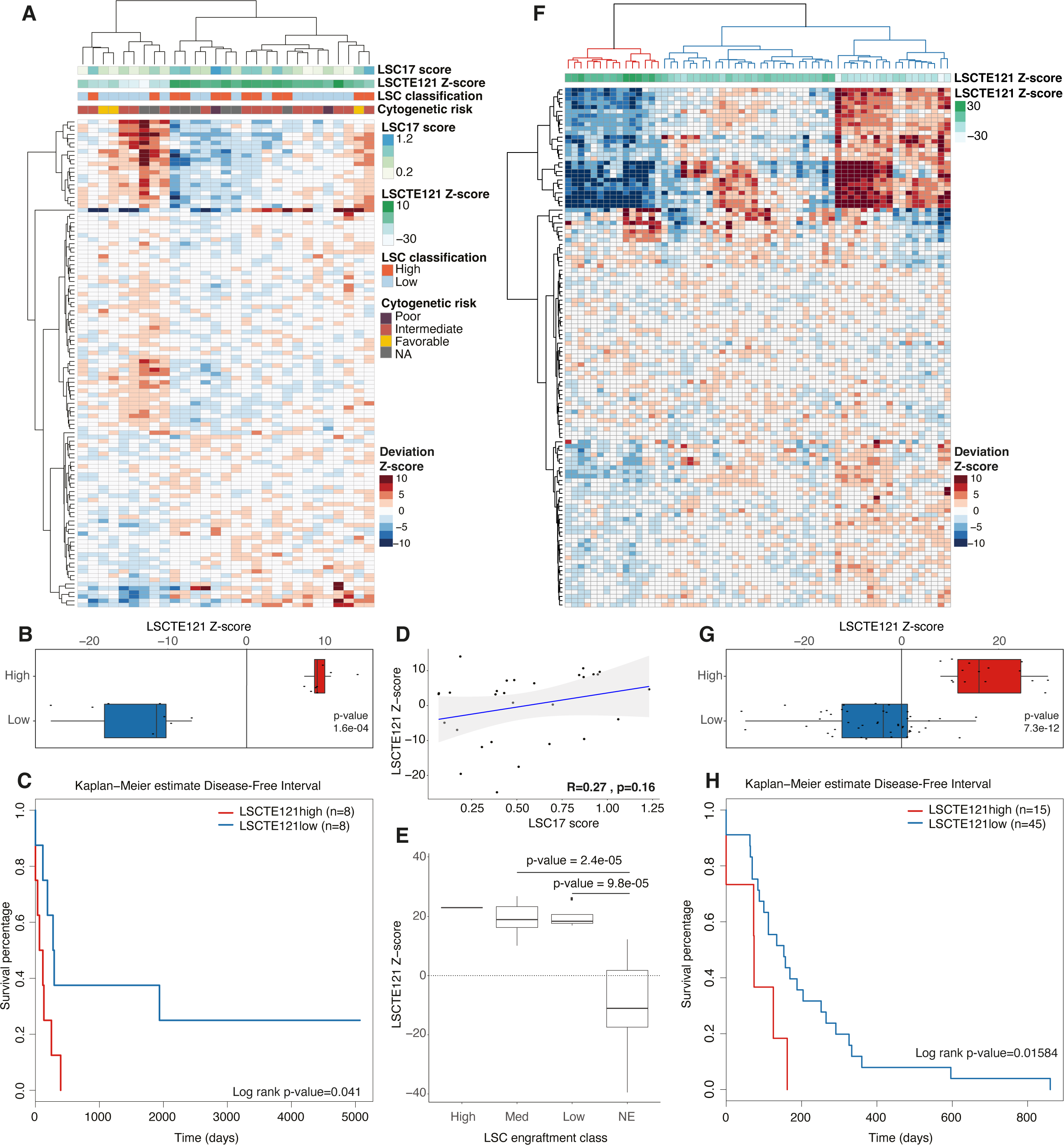
The LSCTE121 signature stratifies AML patients with distinct disease-free intervals. A) Heatmap showing differential accessibility Z-scores for LSCTE121 (q < 0.01 LSC+ vs LSC-) in bulk cohort1 AML tumors. Note that bulk tumors did not cluster according to their LSC17 scores with LSC+ or LSC-fractions. B) Box plot displaying the LSCTE121 Z-score in cohort1 patients. Patients were divided into LSCTE121 high or low based on the heatmaps in A and B. p-values results of the Wilcoxon test are showcased on the box plots. C) Kaplan–Meier estimates using Disease Free Interval as a clinical endpoint for LSCTE121 Z-score top vs bottom 25 percentile in cohort1 (n=8/group). D) Pearson correlation between LSC17 score and LSCTE121 Z-score in cohort1 bulk AML patients. E) Boxplot of LSCTE121 Z-score in LSC fractions. Note that engrafting fractions show higher LSCTE121 Z-score than non-engrafting fractions. F) Same as panel A but for cohort2 patients. G) Same as panel C but for cohort2 patients. H) Same as panel D but for cohort2 patients (LSCTE121high: n=15; LSCTE121low: n=45).

### TE subfamilies accessible in LSCs are docking sites of AML essential factors

Using the ReMap cistrome data, we next assessed if accessible TE subfamilies that were shared or unique across primitive hematopoietic populations and LSC+ fractions (Fig. 2E; Extended Data Tables 10-12) could provide docking sites for different transcription factors. Cistromes for four transcription factors were found to be enriched across accessible TE subfamilies in LSC+ and HSPC populations (Fig. 4A-B and Extended Data Fig. 5A-B). These included ERG, RUNX1, LMO2 and TRIM28. Each of these transcription factors is documented to contribute to normal hematopoiesis and/or leukemia development ^41,50–52^. Beyond its reported role in erythroblast differentiation ^51^, TRIM28 is also known to suppress TE function ^53^, suggesting that it may play a role in the transition from HSPC and LSC+ to mature fates. In agreement with previous studies reporting an interplay between ERG and RUNX1 in hematopoiesis and AML ^54^, the ERG and RUNX1 cistromes were enriched over more than 70% of the same accessible TE subfamilies (10/15 for HSPC-specific, 8/10 for LSC+ specific and 6/8 for TEs shared between HSPC and LSC+ Extended Data Fig. 5C), including LTR78 (ERG: q=0.005, logOR=1.74; RUNX1: q=0.0004, logOR=1.9), LTR67B (ERG: q=1.12e-08, logOR=2.73; RUNX1: q=5.77e-17, logOR=4.09) and MLT1E2 (ERG: q=0.0003, logOR=2.02; RUNX1: q=0.009, logOR=1.73 and Extended Data Fig. 5D, Extended Data Table 10-12). Over half of the TE subfamilies enriched for ERG and/or RUNX1 cistromes also preferentially harbored their DNA recognition sequences (8/25 for ERG motif and 11/32 for RUNX1 motif; Extended Data Fig. 5E-F). Collectively, these results reflect the shared biology across HSPC hematopoietic populations and LSC, relying on four core transcription factors, namely ERG, RUNX1, LMO2, and TRIM28, using accessible elements from TE subfamilies as direct docking sites on the genome.

**Fig. 4:**
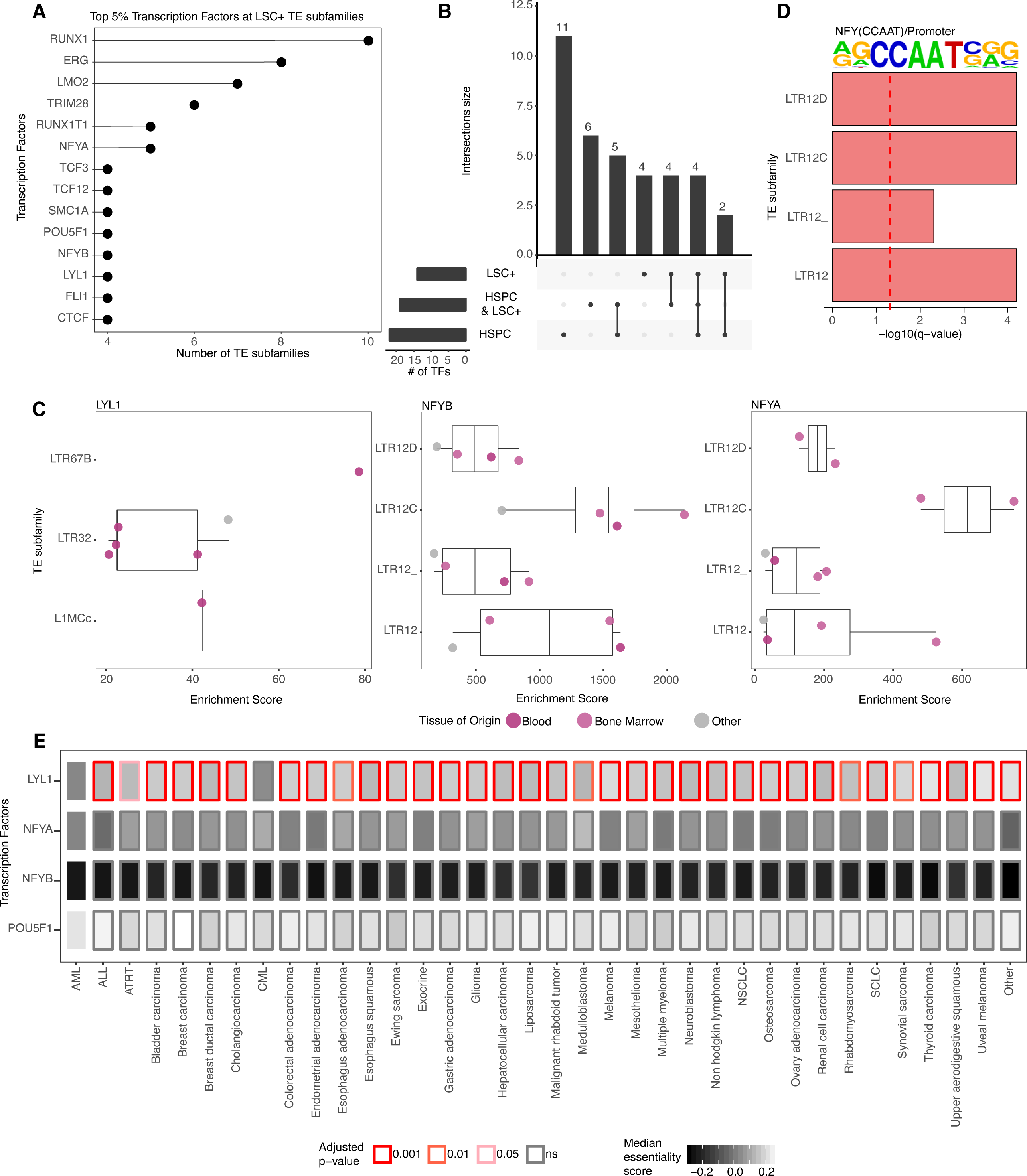
TE subfamilies enriched in accessible chromatin in LSC provide binding sites to transcription factors essential for AML. A) Frequency plot of transcription factor cistromes enriched at TE subfamilies in accessible chromatin in LSC+ fractions. The top 5% most frequently enriched transcription factor cistromes are shown. B) UpSet plot showing the intersection of transcription factor cistromes enriched in LSC+/HSPC populations. C) GIGGLE score for individual LYL1, NFYA, and NFYB cistromes profiled in cell lines derived from various tissue states when compared to TE subfamilies specifically accessible in LSC+. D) Enrichment of NFY DNA recognition sequences within TE subfamilies enriched in the accessible chromatin of LSC+. The red dashed line corresponds to -log10(q)=1.3 (q=0.05) threshold. E) Overview of essentiality scores of LSC+ specific transcription factors across all cancer types available in DepMap with more than five cell lines, based on CRISPR data. The distribution of the essentiality scores of each transcription factor calculated in AML cell lines was compared to the distribution of the same transcriptional factor calculated in cell lines from every other cancer type. Rectangles inner color corresponds to the median essentiality score, while border color corresponds to the pairwise t-test, Benjamini-Hochberg corrected p-values (adjusted p-value).

Cistrome enrichment analysis over accessible TEs identified eleven transcription factors specific to HSPC populations, including HIF1A, SETDB1, GATA1, ZNF143, TEAD4, PBX2, EZH2, CBX3, SMC3, MAX, HDAC2 (Fig. 4B, Extended Data Fig. 5A). While no reports implicate CBX3 and MAX in hematopoiesis, a previous characterization of PBX2 knock-out mice showcased that PBX2 is not essential for normal hematopoiesis or immune response ^55^. Similarly, TEAD4 favors embryonic to blood cell state commitment ^56^, and GATA1, HIF1A, SETB1, EZH2, and HDAC2 are known regulators of HSCs self-renewal and/or differentiation ^57–61^. ZNF143 and SMC3 define a unique class of transcription factors that regulate the three-dimensional genome organization by binding to chromatin interaction anchors in collaboration with CTCF ^62–64^. In HSPC populations, ZNF143, SMC3, CTCF cistromes, and those of their known partners SMC1A, RAD21 and CHD8 ^62,65,66^, enrich over LTR41B elements (q≤3.22e-34, OR≤9.36) (Extended Data Fig. 5D, Extended Data Table 10), in line with the role in 3D chromatin organization of LTR41B elements ^67^. The accessibility of TE subfamilies serving as docking sites for known HSC factors and/or chromatin interaction factors parallels the enrichment of the GATA1 and CTCF DNA recognition sequences at some of these TEs (Extended Data Fig. 5G) and is in keeping with the role of CTCF as a gatekeeper of stemness in hematopoietic populations ^13^. Together, these results suggest that specific TE subfamilies are accessible in HSPC populations, serving as DNA docking sites for HSC factors and/or regulators of genome topology.

Finally, we found four transcription factors whose cistromes were uniquely enriched over accessible TE subfamilies in LSC, namely LYL1, NFYA, NFYB, and POU5F1 (Fig. 4A-B, Extended Data Table 11). From the GIGGLE enrichment score analysis, we observed a bias for this enrichment for cistromes generated in blood or bone marrow samples for LYL1, NFYA and NFYB (Fig. 4C). TE subfamilies associated with NFYA and NFYB binding were also enriched for the DNA recognition sequence for these transcription factors (Fig. 4D). We next assessed the necessity of these transcription factors to growth in AML cell lines. Analyzing the CRISPR-based knock-down essentiality screen results from the Cancer Dependency Map (DepMap) project (see Methods) revealed negative essentiality scores for all four transcription factors across the majority (12/23) of AML cell lines (Extended Data Fig. 6A). While the median essentiality score across all 23 AML cell lines was not significantly lower for POU5F1 compared to all other genes, LYL1, NFYA and NFYB were consistently reported as essential (p-value<0.05; Extended Data Fig. 6A). No significant essentiality for these transcription factors were detected from the DepMap RNAi-based knock-down as opposed to knock-out screen (Extended Data Fig. 6B). When comparing essentiality scores across cells classified by cancer types, LYL1 was significantly more essential in AML cell lines than cells from other cancer types (CRISPR-based screen: 97%, 34/35; RNAi-based screen: 75%, 17/21) (Fig. 4E, Extended Data Fig. 6C, Extended Data Tables 13,14), in agreement with its reported LYL1 pro-oncogenic role in AML ^68^. NFYA and NFYB factors were found essential across all cancer types (Fig. 4E, Extended Data Fig. 6C). Collectively, our results suggest that the accessibility of TE subfamilies specific to LSC defines docking sites for transcription factors, including LYL1, NFYA, and NFYB essential in AML.

### TE subfamilies accessible in LSC are essential for stemness properties

We undertook functional studies to assess the contribution of TEs to stemness properties using the OCI-AML22 model of AML, known to harbor functionally assessed LSC+ and LSC-populations that can be enriched based on cell surface marker expression (CD34+CD38-fraction, very high LSC frequency; CD34+CD38+, very low LSC frequency; and CD34-(either CD38+ or CD38-) fractions, LSC-) ^44^. From ATAC-seq profiles generated across each OCI-AML22 fraction, we measured TE subfamily enrichment within accessible chromatin. Using the LSCTE121 signature for unsupervised clustering, we observed that the LSC+ fraction of OCI-AML22 cells (CD34+/CD38-) clustered with LSC+ AML fractions (Fig. 5A). In contrast, the OCI-AML22 fractions functionally classified as LSC-(CD34-) ^44^ clustered with the majority of LSC-AML fractions (Fig. 5A). This demonstrates that the OCI-AML22 model faithfully recapitulates the heterogeneity in chromatin accessibility found across LSC+ and LSC-populations in primary AML samples, where LSC+ cells drive the production of LSC-populations. In agreement with increased LSC+ content in the CD34+/CD38-versus CD34+/CD38+ fractions in OCI-AML22 ^44^, the LSCTE121 score was higher in the CD34+CD38-compared to CD34+/CD38+ OCI-AML22 fraction (Extended Data Fig. 7A-B). This observation reflects the value of the LSCTE121 score to quantify stemness potential in AML.

**Fig. 5:**
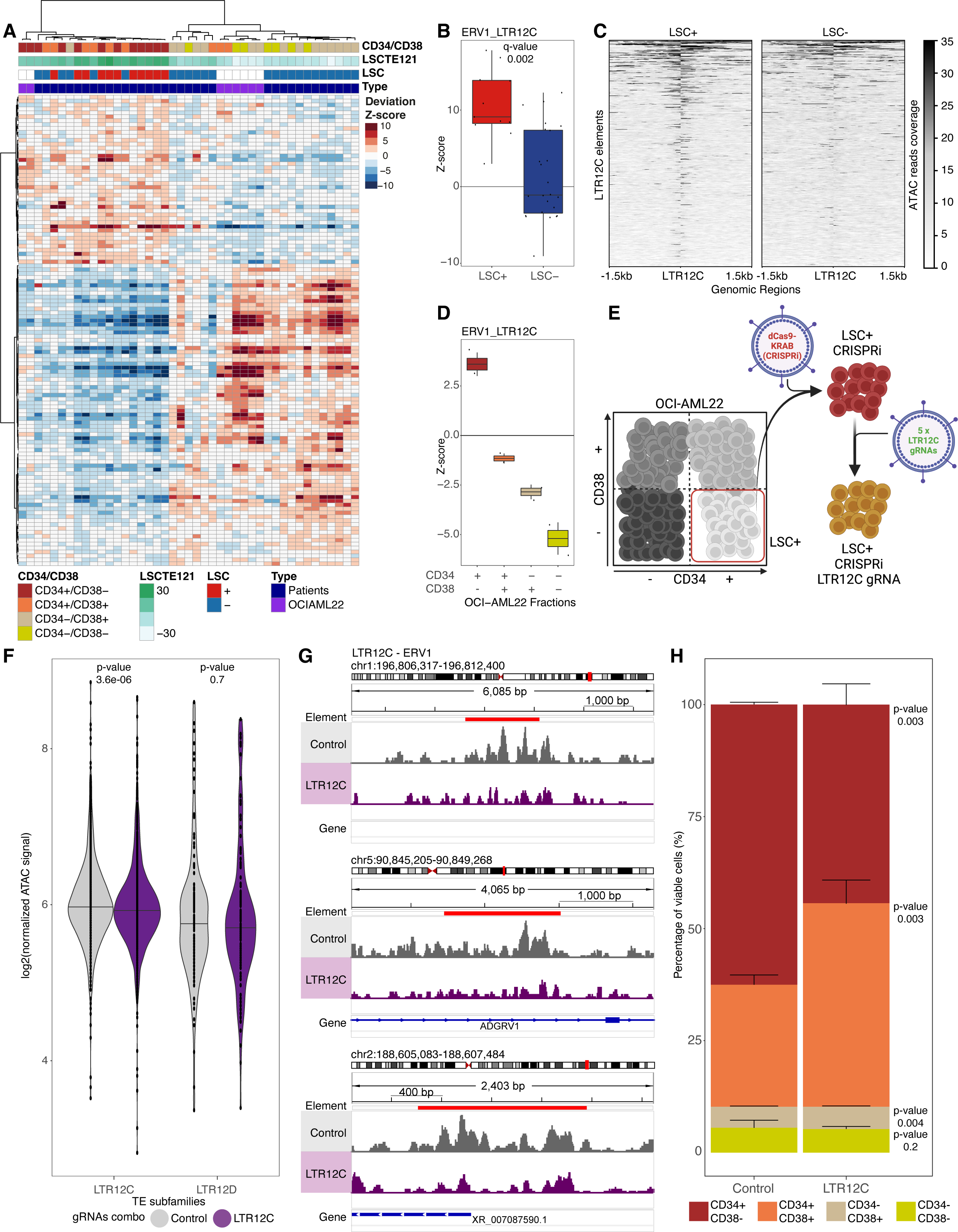
Accessibility at LTR12C elements is essential for LSC stemness properties. A) Heatmap showing differential accessibility Z-scores for LSC TEs (q < 0.01 LSC+ vs LSC-) between ATAC-seq profiles of LSC+, LSC-, OCI-AML22 fractions, OCI-AML3 and MOLM13 cell lines, clustered based on correlation. B) Box plot displaying the LTR12C accessibility Z-scores in LSC+ and LSC-. C) ATAC-seq profile across all LTR12C elements accessible in LSC+ and/or LSC-fractions. D) Same as B but for OCI-AML22 fractions. E) Schematic overview of the CRISPRi chromatin editing approach. F) Violin plot for the chromatin accessibility signal distribution in OCI-AML22-LSC+ stably expressing both CRISPRi and scramble (control in gray) or LTR12C (purple) guide RNA combos over LTR12C or LTR12D elements. A representative biological replicate of three is shown. Every dot represents an individual TE element. p-values results of paired samples Wilcoxon test are showcased on the violin plot. G) Genome browser views of ATAC-seq signal reduction at individual LTR12C elements in OCI-AML22-LSC+ expressing both CRISPRi and LTR12C guide RNA combo (purple) compared to scramble gRNA combo (control in gray). H) Percentage of CD34+/CD38-, CD34+/CD38+, CD34-/CD38+ and CD34-/CD38-cells upon CRISPRi-mediated chromatin editing at LTR12C elements in OCI-AML22-LSC+. The results are shown as means ± SD from three independent transductions with the vector promoting the expression of either 5 scramble guide RNAs or 5 guide RNAs targeting LTR12C elements. p-value was generated by a two-sided t-test.

To directly test whether TE subfamilies are functionally required to drive stemness properties we turned to novel chromatin editing technologies that allow altering the chromatin state across elements from an individual TE subfamily. We focused on the LTR12C TE subfamily, which exhibited one of the most differential chromatin accessibility between LSC+ and HSPC populations (Extended Data Fig. 3C). As expected, the LTR12C subfamily was more enriched in accessible chromatin in LSC+ compared to LSC- and LTR12C elements were overall more accessible in LSC+ compared to LSC-fractions (Fig. 5B-C). Furthermore, the LTR12C subfamily was more enriched in chromatin accessibility in CD34+/CD38-LSC+ compared to other fractions from OCI-AML22, and the enrichment decreased concomitantly as cells lost stemness and became more mature across the cellular hierarchy (Fig. 5D). To functionally assess the role of the LTR12C subfamily towards LSC+ properties, we stably expressed the CRISPR/dCas9-KRAB (CRISPRi) construct ^12,69,70^ in the OCI-AML22 model and evaluated if altering the chromatin state of LTR12C elements could inhibit LSC+ stemness and its ability to regenerate a cellular hierarchy (Fig. 5E, Extended Data Fig. 7C-D, see Methods). By using the Repguide tool ^71^, we generated a combo of 5 guide RNAs targeting specifically the LTR12C subfamily (n=2765 genomic copies), predicted to recognize 2400 copies (87%). As a negative control, we used a combo of 5 scramble guide RNAs (Extended Data Table 15). To test the specificity of our CRISPRi targeting strategy, we profiled chromatin accessibility by ATAC-seq in CRISPRi expressing OCI-AML22 cells co-expressing either scramble or LTR12C guide RNAs. All three biological replicates revealed a decreased chromatin accessibility at LTR12C elements (p≤8e-04) with limited off-target effect when assessed on phylogenetically proximal TE subfamilies, such as LTR12D (p>0.5) (Fig. 5F-G, Extended Data Fig. 7E-G), reflecting the specificity of the chromatin editing approach towards LTR12C elements. Functional flow based analysis showed that when, chromatin was repressed over LTR12C elements there was a reduced percentage of the LSC+ cellular fraction (CD34+/CD38-fraction) as compared to control guides (CD34+/CD38-percentages: 62.5% in control and 44.3% in LTR12C, Fig. 5H). In parallel there was an increase of CD34+/CD38+ cells that represent more committed phenotype (CD34+/CD38+ percentages: 27.2% in control and 45.4% in LTR12C, Fig. 5H). Collectively, these results establish that accessibility of LTR12C elements is a stemness determinant in LSC.

## Discussion

Our study establishes that enrichment of specific TE subfamilies within chromatin variants serves as a hallmark of stemness in both normal hematopoiesis and AML, with direct clinical implications for AML patients. Many of the TE subfamilies enriched in chromatin variants of both LSC and HSPC populations are shared with those found enriched in chromatin variants identified in embryonic stem versus mature cells ^29,72^. This includes members of the endogenous retrovirus (ERV) families known to act as docking sites for stem transcription factors, such as OCT4 and NANOG ^29,36,37^. Expanding on the role of TEs as a source of *cis*-regulatory elements in stem and mature cells ^28,29,36–39,72^, our results show that HSPC-accessible TEs, such as LTR41, LTR41B and LTR55 are permissive for binding by regulators of the three-dimensional genome organization, including ZNF143, CTCF, and cohesin elements. CTCF-driven changes to the genome topology in primitive hematopoietic populations were identified as a regulator of quiescence exit and long-term self-renewal^13^. Here, additional transcription factors, namely ERG, RUNX1, LMO2, and TRIM28, were found based on the presence of their occupancy across elements from the LTR78, LTR67B and MLT1E2 TE subfamilies enriched in chromatin variants from LSC and HSPC populations. ERG, RUNX1, LMO2, and TRIM28 correspond to known hematopoietic transcription factors and reinforce their direct role in AML ^41,50–53^. Collectively, our results suggest a contribution of distinct TE subfamilies to serve as determinants of stem properties, shaping both genome topology and transcriptional regulation and defining the cell state hierarchy in both normal and disease settings.

By revealing how differentially enriched TE subfamilies distinguish LSCs from mature leukemic cells, we provide a chromatin-based approach to interrogate repeat DNA sequences and discriminate AML samples and/or isolated populations according to their leukemia-initiating potential. When applied to AML patient samples, our LSCTE121 scoring scheme can stratify patients to identify those with high rates of relapse and shorter overall survival. The lack of correlation between our LSCTE121 and the LSC17 ^15^ scoring scheme argues for their complementarity in assessing the LSC content in primary patient samples. Although there are large AML datasets of RNA, there are few that interrogate chromatin variants at scale; our work points to the need for a deeper study of the repetitive genome and its contribution towards stemness properties in leukemia, but also cancer in general.

In cancer, chromatin variants are predominantly found at TEs ^73^ and can lead to oncogene activation ^31^. However, studies on the role of TEs in cancer have primarily focused on their expression. For instance, high levels of ERV1 expression are linked to worse survival in patients with kidney cancer ^74^, and a 14 TE-based signature predicts survival in AML ^75^. Our work extends beyond this TE expression-based perspective to the role of TEs as regulatory elements. Among the TE subfamilies accessible in LSCs, for example, we observed the LTR12C, whose elements work as enhancers in AML ^33^ and were previously shown to act as alternative promoters in hepatocellular carcinoma ^76^. We further show that accessible TEs provide docking sites for oncogenic transcription factors in LSCs. Using this approach, we identified LYL1 as an oncogenic driver of AML, in agreement with previous reports ^68^. We further showed that, unlike NFYA/B factors that were essential across all cancer states, LYL1 is preferentially essential across AML cell lines as compared to cell lines from other cancer states. Altogether, our work suggests a direct contribution of the repetitive genome towards the governance of oncogenic transcription factors in AML.

Previous studies reported how altering the chromatin state at TEs with CRISPRi approaches could impact tumor growth in AML and prostate cancer ^29,33^. We used the same technology to showcase the essential role of LSC+-specific TE subfamilies in maintaining stemness. We were specifically able to target hundreds of LTR12C elements, promoting their repression at the chromatin level without altering the underlying sequence. Such chromatin repression caused a drop in the percentage of LSC+ cells and an increase in more mature cell types in OCI-AML22 cells. These results reveal the essential role of LSC+-specific TEs on stemness properties and uncover LSCs’ genetic vulnerabilities that, when targeted, should not affect HSPC populations. Altogether, the work presented here provides a framework to capture the primitive nature of cancers by revealing the role of TE subfamilies as genetic determinants of stemness properties in normal and leukemic stem populations.

## Methods

### Patient samples

All biological samples were collected with informed consent according to procedures approved by the Research Ethics Board of the University Health Network (UHN; REB# 01-0573-C) and viably frozen in the Leukemia Tissue Bank at Princess Margaret Cancer Centre/ University Health Network. Patient details can be obtained from Extended Data Table 9.

### Cell sorting and Xenotransplantation assay

Hematopoietic populations were selected using the highest purity sorting scheme available ^77,78^. Peripheral blood AML samples were sorted into fractions based on the expression of CD34 and CD38. LSC activity in each fraction was assessed by xenotransplantation into *NOD. Prkdc^scid^. Il2rg ^null^*(NSG) mice as previously described ^15^. Antibodies used are shown in Extended Data Table 16.

### Bulk ATAC-seq

ATAC-seq was used to profile the accessible chromatin landscape of 11 LSC+, 24 LSC-fractions, and two independent cohorts of peripheral blood AML bulk samples (cohort1: n=29 12 LSC17 high, 17 LSC17 low; cohort2: n=60, unknown LSC17 score). 50,000 cells from each sample were processed as previously described ^79^. Libraries were sequenced with 50bp single-end (LSC fractions) or 50bp paired-end (AML bulk samples) reads and mapped to hg38. Reads were filtered to remove duplicates, unmapped or poor quality (Q < 30) reads, mitochondrial reads, ChrY reads, and those overlapping the ENCODE blacklist. Following alignment, accessible chromatin regions/peaks were called using MACS2. Default parameters were used except for the following: --keep-dup all -B --nomodel --SPMR - q 0.05. ATAC-seq profiles of hematopoietic cell populations were downloaded from GSE125064 and processed similarly. We only included LSC fractions from patients without NPM1 mutation and set a threshold of at least 10 million reads and more than 15k called peaks. ATAC-seq data of LSC fractions and bulk AML samples can be accessed at EGAS00001004893, EGAS00001004896, and EGAS00001007191, respectively. Narrowpeak files are submitted to GEO under ID GSE234237.

### TE families and families

Bedfiles of repeats classified as TE families (excluding hence simple repeats, satellites, sn/rRNA, and repeats of unknown families) and with more than 100 individual genomic elements (n=971) in the hg38 human genome build were downloaded from the UCSC Genome Browser. Subfamilies were classified into families according to Repbase (https://www.girinst.org/repbase/) ^80,81^.

### TE enrichment in accessible chromatin analysis

TE enrichment analysis was performed using ChromVAR ^82^. The binary matrix for ChromVAR was generated by identifying the presence/absence of accessible chromatin peaks in each sample of interest. For the same set of peaks, a binary annotation matrix was generated with rows as peaks and columns as each TE subfamily. TE subfamily-to-peak assignment was generated by checking the overlap of TEs for each subfamily in the accessible chromatin peak set. ChromVAR was run with default parameters using the two input matrices, generating a bias-corrected deviation Z-score. Deviation Z-scores were then compared between samples of interest using a non-parametric two-sided Wilcoxon signed rank test. For plotting, heatmap colors for Z-scores were truncated to minimum and maximum values of -10 and 10, respectively, and clustering was based on Euclidean distance unless otherwise stated. For the LSC TE Z-score, we combined all LSC+/LSC- enriched TEs in separate .bed files and calculated the Z-score over all LSC+/LSC- enriched TEs separately. To combine both Z-scores, we used Stouffer’s method with equal weights of the LSC+ Z-score and the inverse LSC- Z-score:

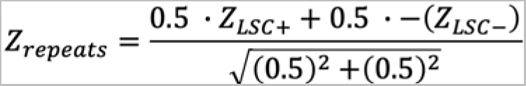

ATAC signal was quantified over LTR12C elements accessible in LSC+ and/or LSC- using deeptools^83^. Given the different number of samples for LSC+ and LSC- fractions (11 and 24 respectively), we reported ATAC signal of top and bottom 25% of samples based on LSCTE121 score. The top 25% corresponds to LSC+, while the bottom 25% is composed uniquely of LSC- and it was therefore named LSC-.

### Transcription factor cistromes and motifs enrichment analysis

The enrichment of cistromes and motifs was assessed at the TE subfamilies enriched specifically in LT-HSCs, HSPC populations, LSC+ and commonly enriched in primitive populations and LSC+(5, 70, 21 and 23 TE families respectively; Extended Data Fig. 2C, Fig. 2E). Each TE subfamily was analyzed individually, restricting our analysis to the elements overlapping with a given consensus set of ATAC peaks (e.g. for a subfamily enriched in primitive populations, we inquired transcription factor cistromes and motifs enriched at the elements overlapping with the ATAC peaks catalog of primitive populations). Enrichment of transcription factor cistromes was performed using the Bioconductor package LOLA (https://bioconductor.org/packages/release/bioc/html/LOLA.html) ^84^, version 1.20.0. 485 transcription factor cistromes were obtained from ReMap (http://remap.univ-amu.fr/) ^85^. Enriched transcription factor cistromes at individual TE subfamilies were called against all elements from the 971 TE subfamilies overlapping any of the primitive/mature/LSC+/LSC- ATAC peaks (q < 0.05). The top 5% most frequently enriched are shown in Fig. 4A, Extended Data Fig. 5A-B. HOMER motif discovery tool (findMotifsGenome.pl) version 4.7 was used to compute the enrichment of motifs. Enriched motifs within individual TE subfamilies (as explained above) were called against all elements from the 971 TE subfamilies overlapping any of the primitive/mature/LSC+/LSC- ATAC peaks (Benjamini q < 0.05). GIGGLE enrichment scores were calculated as described in ^29^ with the following parameters: Species - Human hg38, Data type in Cistrome - Transcription factor, chromatin regulator, Peak number of Cistrome sample to use - All peaks in each sample.

### Essentiality scores

Essentiality scores were obtained from the Broad Institute project Achilles (DepMap) ^86^. CRISPR (Avana 21Q1) and combined RNAi genetic dependencies data were downloaded from the “Download” page of the website https://depmap.org/portal/achilles/. For CRISPR data, we classified cell lines into cancer types following the sample information available on the DepMap website. In particular, the following 23 cell lines were classified as AML: HEL, HEL 92.1.7, MV411, OCI-AML2, THP1, NOMO-1, EOL-1, KASUMI-1, N-4, OCIAML3, MOLM13, TF1, U937, F36P, KO52, AML-193, M-07e, P31/FUJ, MONO-MAC-1, MOLM14, MUTZ-8, OCI-M2 and SHI-1. For RNAi data, our analysis was restricted to the following AML cell lines analyzed by the project Achilles: HL60, MOLM13, MOLM16, MV411, NB4, OCIAML2, OCIAML3, OCIAML5, THP1 and U937.

### LSC17 scores

The LSC17 scores for all AML bulk samples were obtained using the clinical LSC17 assay. In general, the LSC17 score can be calculated as previously reported ^15^. In short, log2 normalized values of each of the 17 genes were multiplied by its coefficient and then summed. For genes with more than one probe, probes with the highest overall signal were selected.

### Survival analysis

The disease-free interval was defined as the time from remission until the date of the first new tumor progression event (locoregional recurrence, distant metastasis, development of a new primary tumor, or death). Survival analysis was performed using the Kaplan–Meier estimate method. P values comparing Kaplan–Meier survival curves were calculated using the log-rank (Mantel–Cox) test.

### OCI-AML22 culturing

Cells were expanded in culture in the following medium, with final concentrations as indicated: X-VIVO 10 (Lonza, BE04-380Q) supplemented with 20% BIT 9500 Serum Substitute (StemCell technologies, 09500), 1x Glutamax Supplement (Thermo Fisher Scientific, 35050061), Primocin 0.1 mg/ml (invivogen), SCF (200 ng/ml; Miltenyi Biotech, 130-096-696), IL3 (20 ng/ml; Miltenyi Biotech, 130-095-069), TPO (20 ng/ml; Peprotech, 300-18), FLT3L (40 ng/ml; Peprotech, 300-19 B), IL6 (10 ng/ml; Miltenyi Biotech, 130-093-934), G-CSF (10 ng/ml; Miltenyi Biotech, 130-093-861). Cells were maintained at a density of 0.8x10^6^ cells/ml and passaged every 3-5 days in a 96-well flat bottom plate. The CD34+CD38- fraction or CD34+ fraction was regularly sorted to serially expand the cells.

### Lentiviral production

VSV-G pseudotyped lentiviral vector particles were produced by polyethyleneimine (PEI)- based co-transfection^87^ of 10.5 μg of pVSVg, 20.5 μg of pCMVR8.74 (both from Addgene) and 38 μg of transfer vector (pHR-SFFV-KRAB-dCas9-P2A-mCherry, from addgene, Plasmid #60954) per HEK293FT cells T175 flask. Viral particles harvested after 44 and 70 hrs were pooled and filtered (0.22-µm filter), concentrated 100xLJusing Lenti-X concentrator protocol (Takaka, ref 631232) optimized with Lenti-X concentrator beads at a ratio 1/8 (beads/virus supernatant). Concentrated virus were resuspended in X-VIVO 10 (Lonza) and stored at –80LJ°C until use.

### Lentivirus titration

Virus titration was assessed on MOLM13 cells. For titration, MOLM13 cells were plated at 25,000 cells/well of 96 well round bottom plate, in 100 μl/well. Viral supernatant was added in 3 serial dilutions in duplicate. After mixing by pipetting cultures were incubated O/N. On the next day, 60-80 μl of supernatant was carefully removed, and 150 μl fresh medium was added. After 3 additional days transduction efficiency (%BFP+) was determined with a flow cytometer (Symphony, BD). Titers were calculated by using the mean of all 6 transductions of duplicate serial dilutions considering poisson-distribution: TU/mL= ln(%untransduced cells/100)* 25,000 cells/(vector input eg. 0.5 μl/1000).

### Generation of dCas9-KRAB (CRISPRi) OCI-AML22 and transduction with LTR12C or control vectors

After expansion, the OCI-AML22 CD34+CD38- fraction was sorted (Moflo, Astrio, BD), washed, and resuspended in its own media. Cells were mixed with concentrated virus produced as previously described, at a 1/10 ratio supplemented with LentiBOOST research grade, Mayflower Biosciences, ref SB-P-LV-101-03 (1/100). Cells were then spun at 1200 rpm for 10 min before leaving them overnight in the incubator. The following day, the medium was replaced with OCI-AML22 culturing medium. To obtain enough cells, OCI-AML22 transduced cells were expanded over 1.5 months before being sorted for the mcherry+CD34+ population. After 2 additional months of cell expansion, mcherry+CD34+ were sorted again (Moflo, BD) to obtain 82 million cells. 1.79 million OCI-AML22 CD34+mcherry+ cells were collected and used for the next steps and are referred to as the dcas9-KRAB (CRISPRi) OCI-AML22. The mcherry+CD34+ sorted cells were transduced the day following sort, with 3 independently produced virus (4 wells for each condition) in a 96-well plate. Each well received 160 000 OCI-AML22 mcherry+CD34+ cells, the virus of interest concentrated as described in the virus section (LTR12C or control) at a dilution 1/10 and LentiBOOST research grade, Mayflower Biosciences, ref SB-P-LV-101-03 (1/100). Cells were then spun at 1200 rpm for 10 min before leaving them overnight in the incubator. The following day, the medium containing virus was washed and replaced with OCI-AML22 culturing medium. 5 days after transduction with LTR12C or control vector guides lentivirus, cell surface marker, mcherry, viability, and GFP expression were assessed on the Symphony, BD using CD34-APC (BD, clone 8G12, ref 340441, 1/100), CD38-PeCy7 (BD, clone Hb7, ref:335790 1/100). 7 days post-transduction, the mcherry+GFP+ population was sorted (MoFlow, BD) for each of the replicate conditions. Enough cells were collected to perform on each replicate ATAC-Seq.

### Design of guide RNAs targeting TEs

Guide RNAs targeting LTR12C elements were designed using Repguide (https://tanaylab.github.io/repguide/; ^88^, using default parameters. We selected the combination of 5 guide RNAs targeting the highest number of LTR12C elements with limited targeting of other LTR12 families (LTR12, LTR12_, LTR12B, LTR12D, LTR12E and LTR12F). Five negative control guide RNAs ^71^ were used as negative control guide RNAs combination. To perform our experiments guide RNAs were cloned in a lentiviral vector (VectorBuilder.com), carrying also the gene coding for the GFP. Sequences of guide RNAs used in this study can be found in Extended Data Table 15.

### Protein extraction and western blot analysis

OCI-AML22 cells stably expressing dCas9-KRAB (CRISPRi) were lysed in modified RIPA (10 mmol/L Tris-HCl, pH 8.0; 1 mmol/L EDTA; 140 mmol/L NaCl; 1% Triton X-100; 0.1% SDS; 0.1% sodium deoxycholate) containing protease inhibitor (cOmplete, EDTA-free Protease Inhibitor combination, Roche). Lysates were sonicated for 5 minutes (30s ON, 30s OFF) using a Diagenode Bioruptor 300. Cell debris was removed by centrifugation (10min, 15,000 rpm at 4°C) followed by protein quantification using the BCA Protein Assay Kit (Thermo Fisher Scientific). Western blot was performed by using the Protein Simple Automated system (Jess, Bio-Techne) following manufacturer instructions. We used an anti-Cas9 antibody (Diagenode, C15200203 - 1:5000) and anti-GAPDH (Cell Signaling, GAPDH (14C10) Rabbit mAb #2118 - 1:300).

## Research reproducibility and code availability

Code for data processing, analysis, and plotting can be found on CodeOcean (https://codeocean.com/capsule/4918318/tree). The analytical pipeline is also made reusable on the CoBE platform (www.pmcobe.ca).

## Authors Contributions

G.G., B.N.., A.Q. and M.L. designed the studies. G.G. and B.N. performed the computational analysis with help from A.Q. A.Mi. performed the functional xenograft transplantation while C.A. and N.T. prepared ATAC-seq libraries. A.Mu. processed part of ATAC-seq data and S.A.M.T gave computational input. Ö.D. and H.B. performed experimental validations. A.A. provided clinical annotation for AML samples. M.D.M provided AML samples. G.G, B.N., A.Q., Ö.D., H.B., J.C.Y.W., M.D.M., J.E.D., M.L. interpreted the data; M.L. and J.E.D. supervised the study. G.G., B.N. and M.L. wrote the manuscript.

## Acknowledgements

We acknowledge the Princess Margaret Genomics and Bioinformatics group for providing the infrastructure required to conduct analyses included in this work. This work was funded by grants to M.L. and J.D. from the Princess Margaret Cancer Foundation, Ontario Institute for Cancer Research with funding from the Province of Ontario, Canadian Institutes for Health Research (CIHR), CEEHRC team grant from the CIHR, Medicine by Design, the Canadian Cancer Society Research Institute; to J.D. from the Terry Fox Research Institute Program Project and a Canada Research Chair; to N.T. from JSPS KAKENHI Grant Number JP17K09899. M.L. holds an Investigator Award from the Ontario Institute for Cancer Research and a Bernard and Francine Dorval Award from the Canadian Cancer Society.

## Supplementary information

Extended Data Table 1. Differential accessible TEs in HSPC vs mature hematopoietic populations

Extended Data Table 2. Differential accessible TEs in LT-HSCs vs ST-HSCs hematopoietic populations

Extended Data Table 3. Differential accessible TEs in stem vs mature hematopoietic populations

Extended Data Table 4. Differential accessible TEs in progenitor vs mature hematopoietic populations

Extended Data Table 5. Differential accessible TEs in stem vs progenitor hematopoietic populations

Extended Data Table 6. Differential accessible TE Zscores in hematopoietic signatures

Extended Data Table 7. Transcription regulator cistromes enriched at stem TEs

Extended Data Table 8. Differential accessible TEs in LSC+ vs LSC-populations

Extended Data Table 9. Clinical information AML patients

Extended Data Table 10. Top 5% transcription regulator cistromes enriched at HSPC populations TEs

Extended Data Table 11. Top 5% transcription regulator cistromes enriched at LSC+ TEs

Extended Data Table 12. Top 5% transcription regulator cistromes enriched at HSPC and LSC+ TEs

Extended Data Table 13. CRISPR-based essentiality scores of LSC+ specific transcription regulators

Extended Data Table 14. RNAi-based essentiality scores of LSC+ specific transcription regulators

Extended Data Table 15. List of guide RNA sequences used in this study

Extended Data Table 16. Antibodies used for FACS sorting of CD34/CD38 fractions

**Extended Data Fig. 1:**
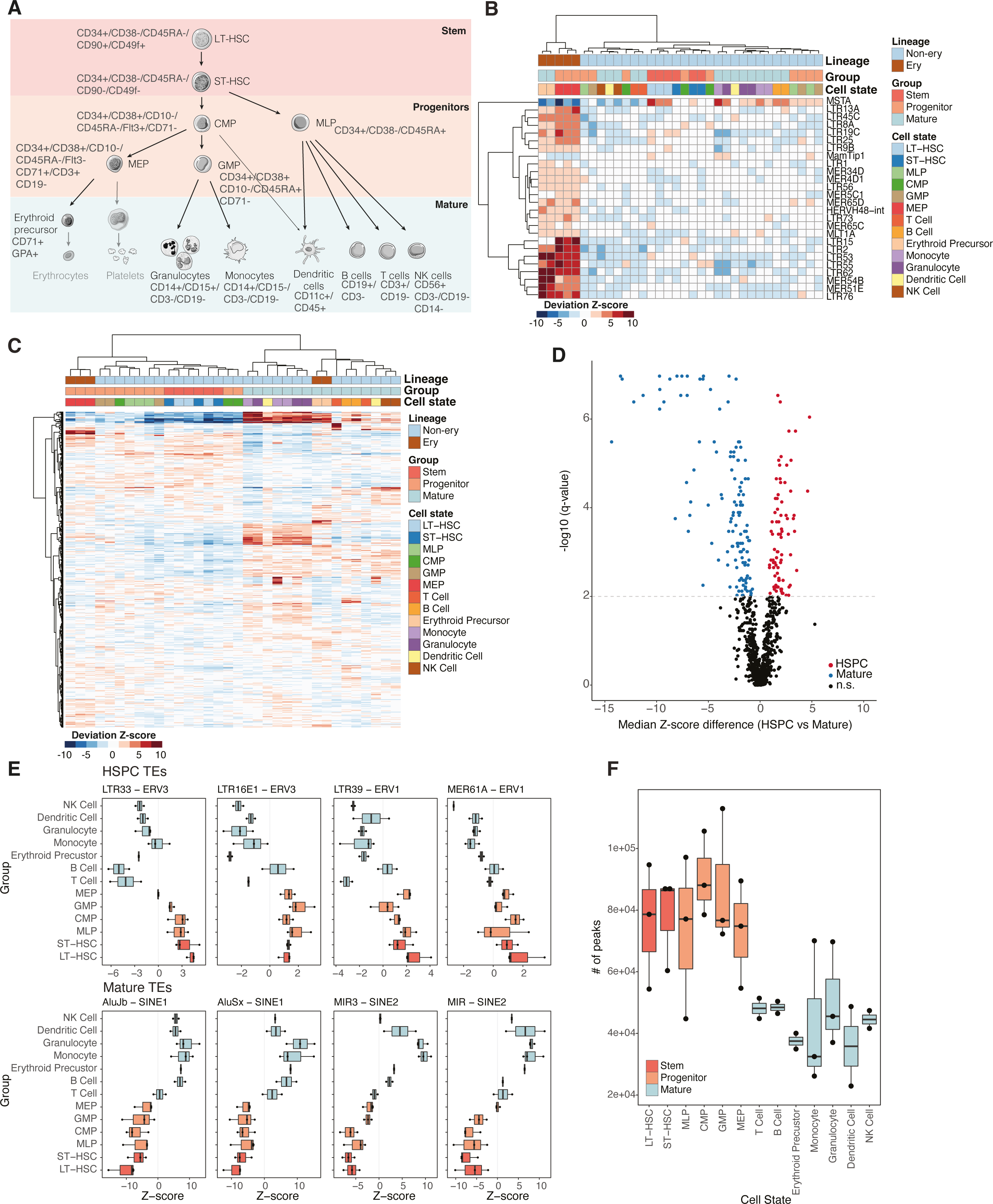
A) Schematic representation of hematopoiesis. B) Heatmap of differential accessibility Z-scores of TE subfamilies with q < 0.01 across bulk ATAC-seq profiles comparing megakaryocyte-erythroid lineage and all other hematopoietic samples. C) Heatmap displaying differential accessibility Z-scores of all TE subfamilies across bulk ATAC-seq profiles of HSCs, progenitors and mature hematopoietic populations excluding megakaryocyte-erythroid lineage TEs. D) Volcano plot showing the median difference in accessibility Z-scores for each TE subfamily between the stem/progenitor and differentiated hematopoietic cell populations versus the −log10 q-value for that difference. E) Examples of non-stem and stem-enriched families of the ERV1, ERV3, SINE1 and SINE2 TE families. Boxplots are showing differential accessibility Z-scores in the different hematopoietic populations. F) Number of called peaks in the different hematopoietic cell populations, note that stem and progenitor cells have a higher number of called peaks than mature hematopoietic cells.

**Extended Data Fig. 2:**
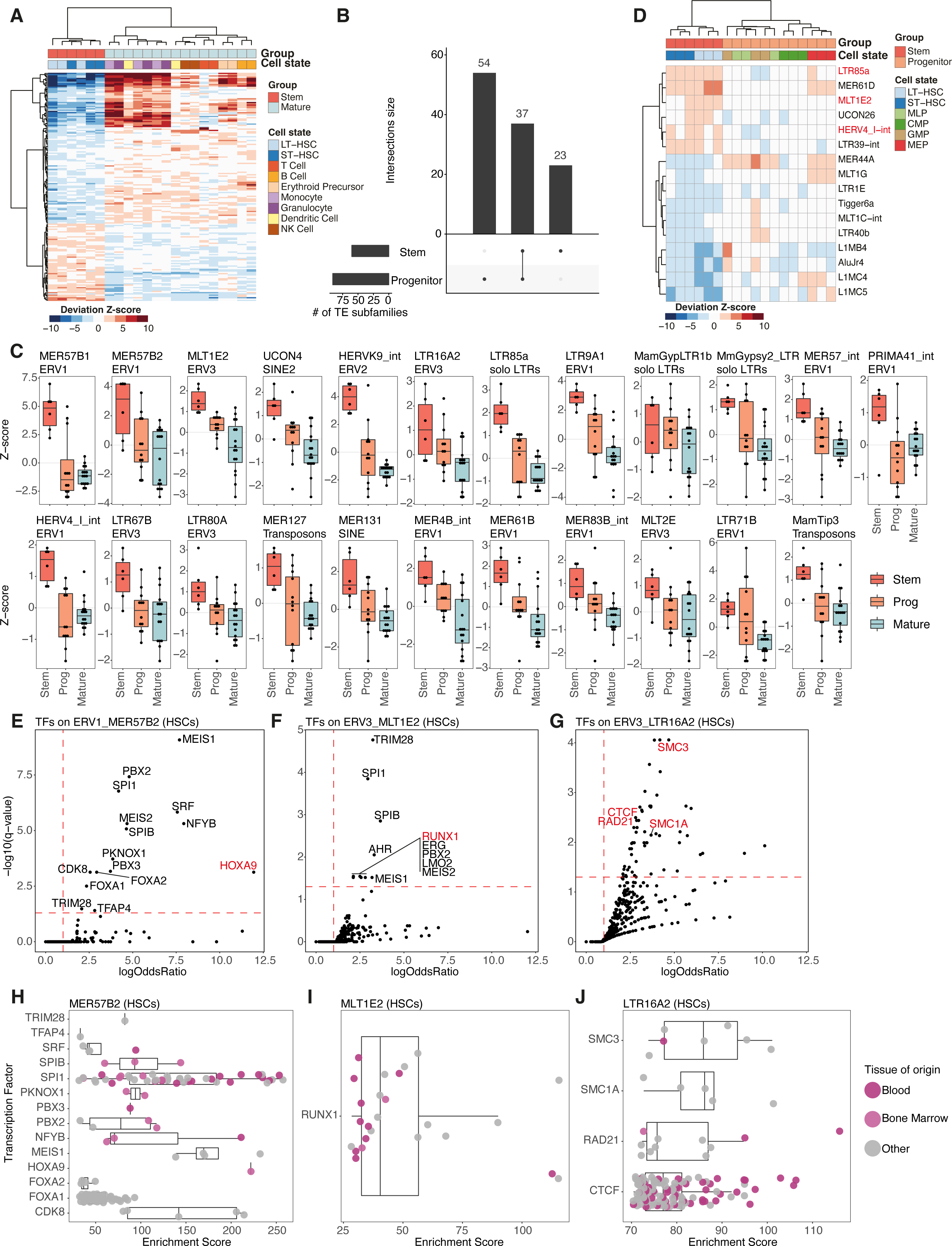
A) Heatmap of differential accessibility Z-scores of TE subfamilies with q < 0.01 across bulk ATAC-seq profiles comparing stem and all mature hematopoietic samples. B) UpSet plot showing the intersection of TE subfamilies enriched in stem or progenitor hematopoietic populations, note that 23 TE subfamilies are enriched exclusively in stem populations C) 23 TE subfamilies enriched exclusively in stem samples. Bar plots show differential accessibility Z-scores in the stem, progenitor and mature cell state groups. D) Heatmap of differential accessibility Z-scores of TE subfamilies with q < 0.05 across bulk ATAC-seq profiles comparing stem and progenitor hematopoietic samples. E) , F) and G) Transcription factor cistromes enriched at 3 of the 23 TE subfamilies enriched in stem populations (HSCs). Every dot corresponds to a transcription factor cistrome taken into account in the analysis. Red dashed lines correspond to -log10(q)=1.3 (q=0.05) and logOddsRatio(logOR)=1 thresholds. H) , I) and J) Individual HOXA9 (H), RUNX1 (I) or three-dimensional genome organization factors (J) cistromes enriched over HSCs specific TE subfamilies and with enrichment of the HOXA9, RUNX1 or three-dimensional genome organization factors cistrome (according to E, F and G). Box plots showing enrichment GIGGLE scores of individual cistromes profiled in cell lines derived from a variety of tissue states.

**Extended Data Fig. 3:**
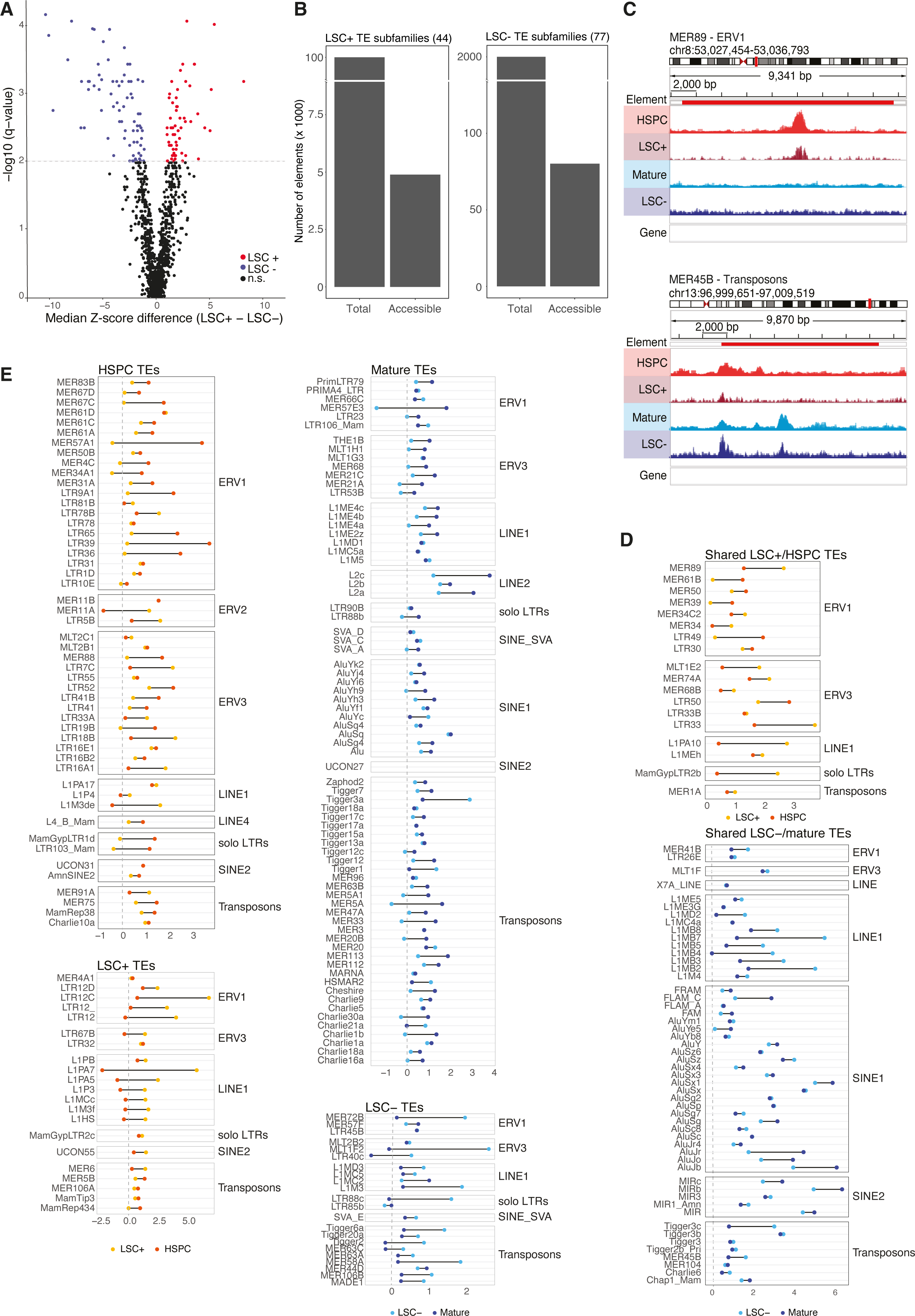
A) Volcano plot showing the median difference in bias-corrected accessibility deviation Z-scores for each TE subfamily between the LSC+ and LSC-fractions versus the −log10(q-value) for that difference. B) Bar plots showing the number of total or accessible TE elements belonging to the 44 TE subfamilies enriched in LSC+ fractions (left) or in the 77 TE subfamilies enriched in LSC-fractions (right). C) Genome browser views of ATAC-seq signal reduction at individual TE elements in HSPC, LSC+, Mature and LSC-samples. D) TE subfamilies are differentially enriched only in LSC+, LSC-, Stem/Progenitor and Differentiated populations of the hematopoietic system, as shown in Fig. 2e. The Cleveland dot plot shows the median z-scores for the relevant comparisons. E) Shared TE subfamilies between LSC+, LSC-, Stem/Progenitor and Differentiated populations of the hematopoietic system as shown in Fig. 2E, Cleveland dot plot shows the median Z-scores for the relevant comparisons.

**Extended Data Fig. 4:**
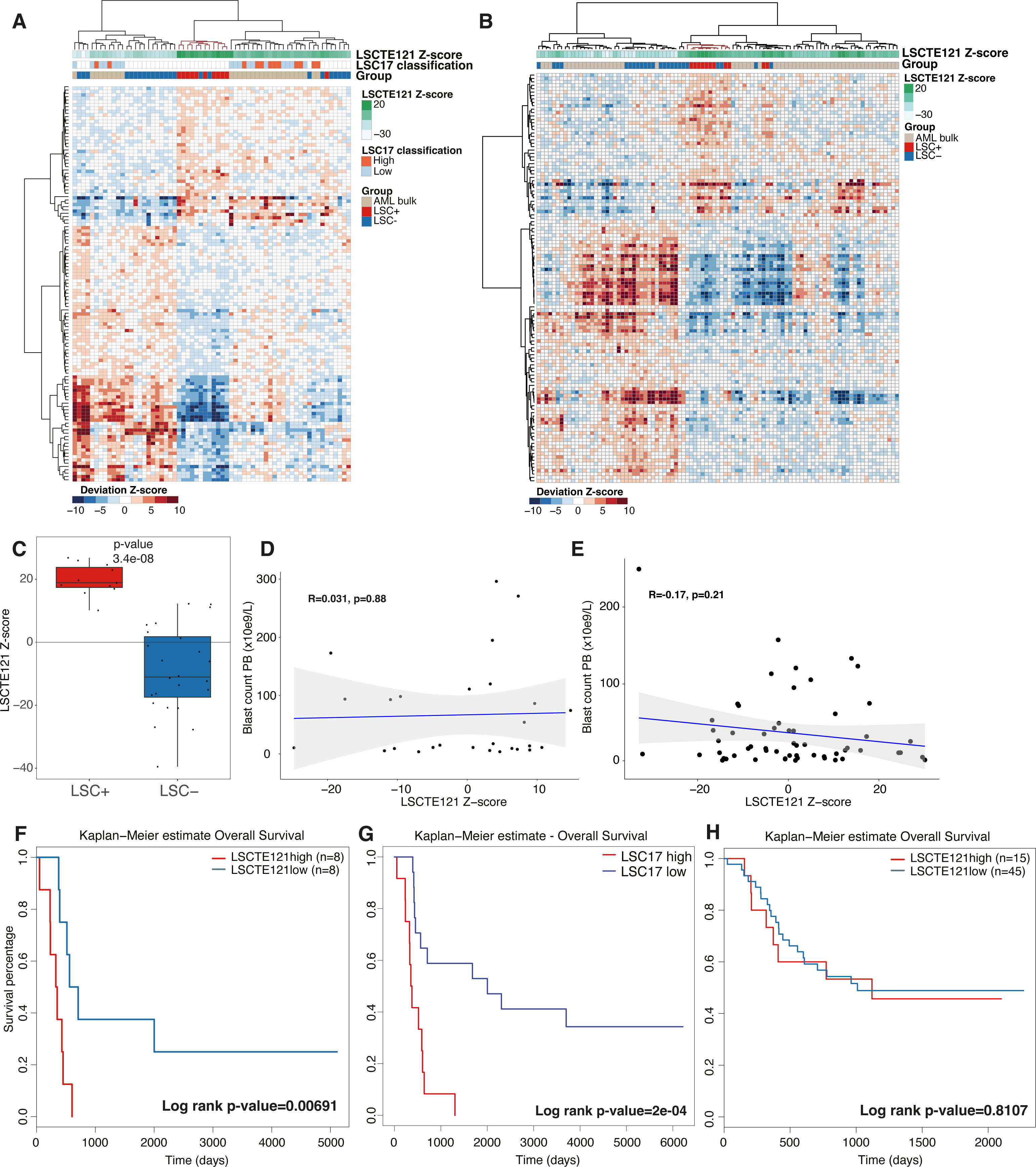
A) and B) Heatmap showing differential accessibility Z-scores for LSC TEs (q < 0.01 LSC+ vs LSC-) between ATAC-seq profiles of LSC+, LSC-fractions and bulk AML tumors (cohort1 in A and cohort2 in B), clustered based on correlation. Note that 20 and 40 samples cluster with LSC+ fractions in cohort1 and cohort2 bulk AML patients respectively. C) Boxplot of LSCTE121 Z-score in LSC fractions, note that engrafting fractions show higher LSCTE121 Z-score than non-engrafting fractions. p values results of the Wilcoxon test are showcased on the box plots. D) Pearson correlation between LSCTE121 Z-score and blast count in peripheral blood for cohort1 bulk AML patients. E) Pearson correlation between LSCTE121 Z-score and blast count in peripheral blood for cohort2 bulk AML patients. F) Kaplan–Meier estimates using Overall Survival as a clinical endpoint for LSCTE121 Z-score top vs bottom 25 percentile in cohort1 (n=8/group). G) Kaplan-Meier estimates using Overall Survival as a clinical endpoint, patient division based on LSC17 high vs low scores in cohort1 bulk AML patients; LSC17 high: n = 12, LSC17 low: n = 17. H) Same as I but for cohort2 patients (LSCTE121high: n=15; LSCTE121low: n=45).

**Extended Data Fig. 5:**
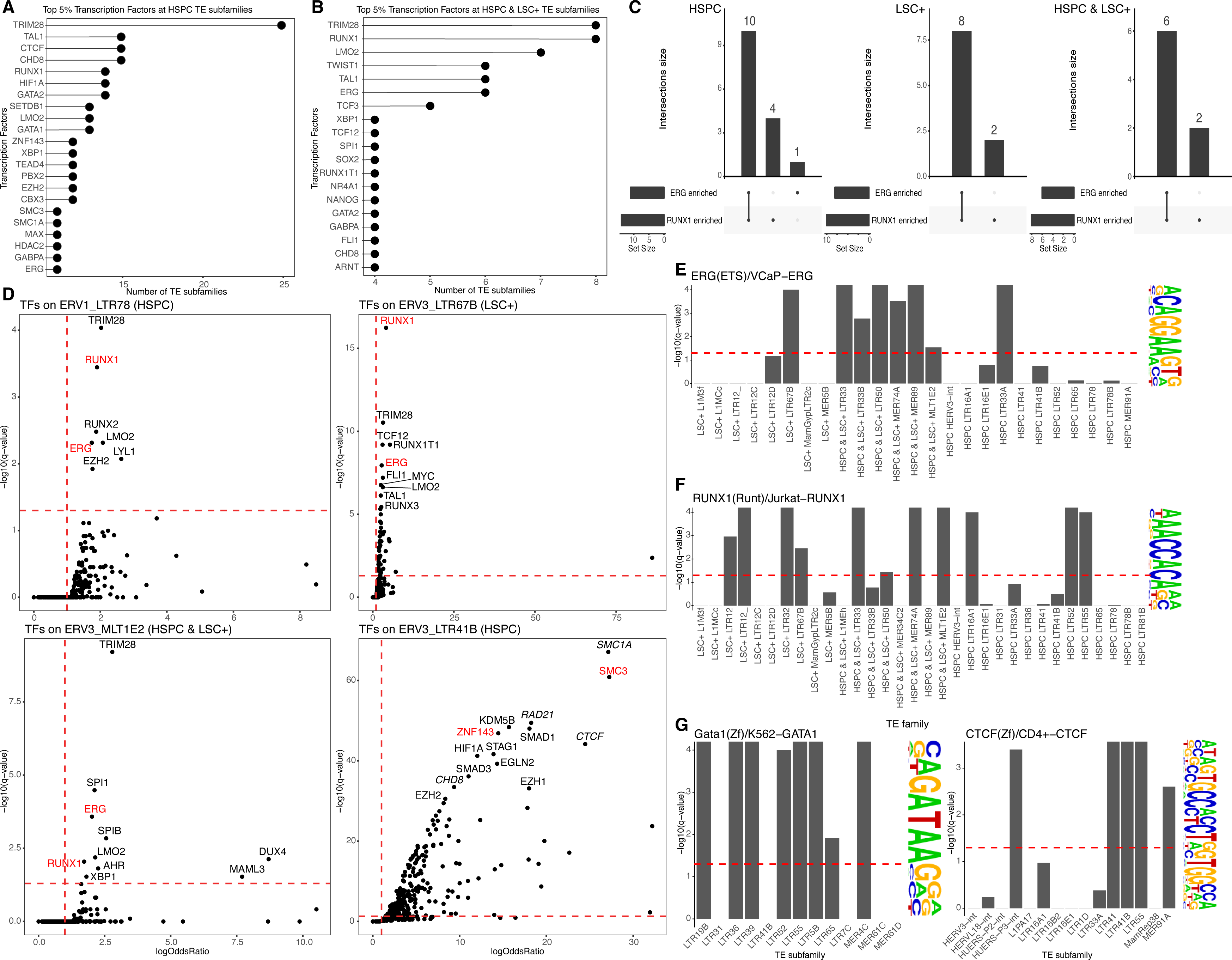
A) Frequency plot of transcription factor cistromes enriched at TE subfamilies enriched in primitive populations. The top 5% most frequently enriched transcription factor cistromes are shown. B) Frequency plot of transcription factor cistromes enriched at TE subfamilies enriched in primitive and LSC+ fractions. The top 5% most frequently enriched transcription factor cistromes are shown. C) UpSet plot showing the intersection of TE subfamilies showing enrichment for ERG and/or RUNX1 cistromes in primitive and/or LSC+. Note that most of the TE subfamilies are shared between ERG and RUNX1. D) Example of TE subfamilies enriched in primitive and/or LSC+ with enrichment of both ERG and RUNX1 cistromes and TE subfamily enriched in primitive populations with enrichment of ZNF143, CTCF and other looping factor cistromes. Dots correspond to each transcription factor cistrome taken into account in the analysis. Red dashed lines correspond to - log10(q)=1.3 (q=0.05) and logOddsRatio(logOR)=1 thresholds. E) Enrichment of ERG DNA recognition sequences within TE subfamilies enriched in LSC+ and/or primitive populations. Bars represent -log10(q). The red dashed line corresponds to - log10(q)=1.3 (q=0.05) threshold. F) Same as E but for the RUNX1 DNA recognition sequences. G) Same as E but for the GATA1 and CTCF DNA recognition sequences within TE subfamilies enriched in primitive populations.

**Extended Data Fig. 6:**
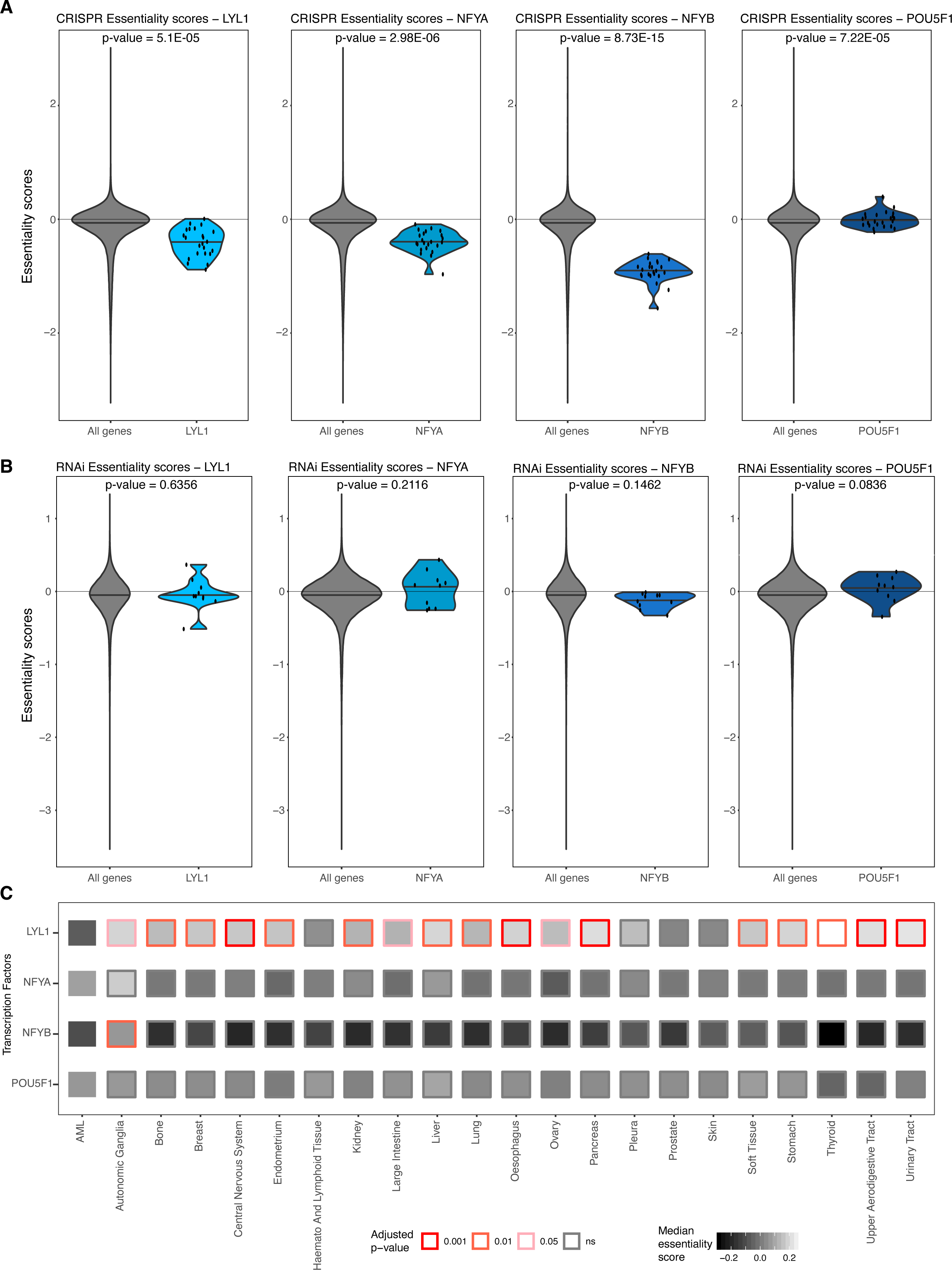
A) Violin plots for essentiality scores of LSC+ specific transcription factors, based on CRISPR data. For each transcription factor the distribution of the essentiality scores calculated in 23 AML cell lines was compared to the distribution of all the other genes tested. p-values based on two-sided t-tests are reported. The red line corresponds to the essentiality score = 0 threshold. B) Violin plots for essentiality scores of LSC+ specific transcription factors, based on RNAi data. For each transcription factor the distribution of the essentiality scores calculated in 10 AML cell lines was compared to the distribution of all the other genes tested. p-values based on two-sided t-tests are reported. The red line corresponds to the essentiality score = 0 threshold. C) Overview of essentiality scores of LSC+ specific transcription factors across all cancer types available in DepMap with more than five cell lines, based on RNAi data. The distribution of the essentiality scores of each transcription factor calculated in AML cell lines was compared to the distribution of the same transcriptional factor calculated in cell lines from each other cancer type. Rectangles inner color corresponds to the median essentiality score, while border color corresponds to Pairwise t-test Benjamini-Hochberg corrected p-values (adjusted p-value).

**Extended Data Fig. 7:**
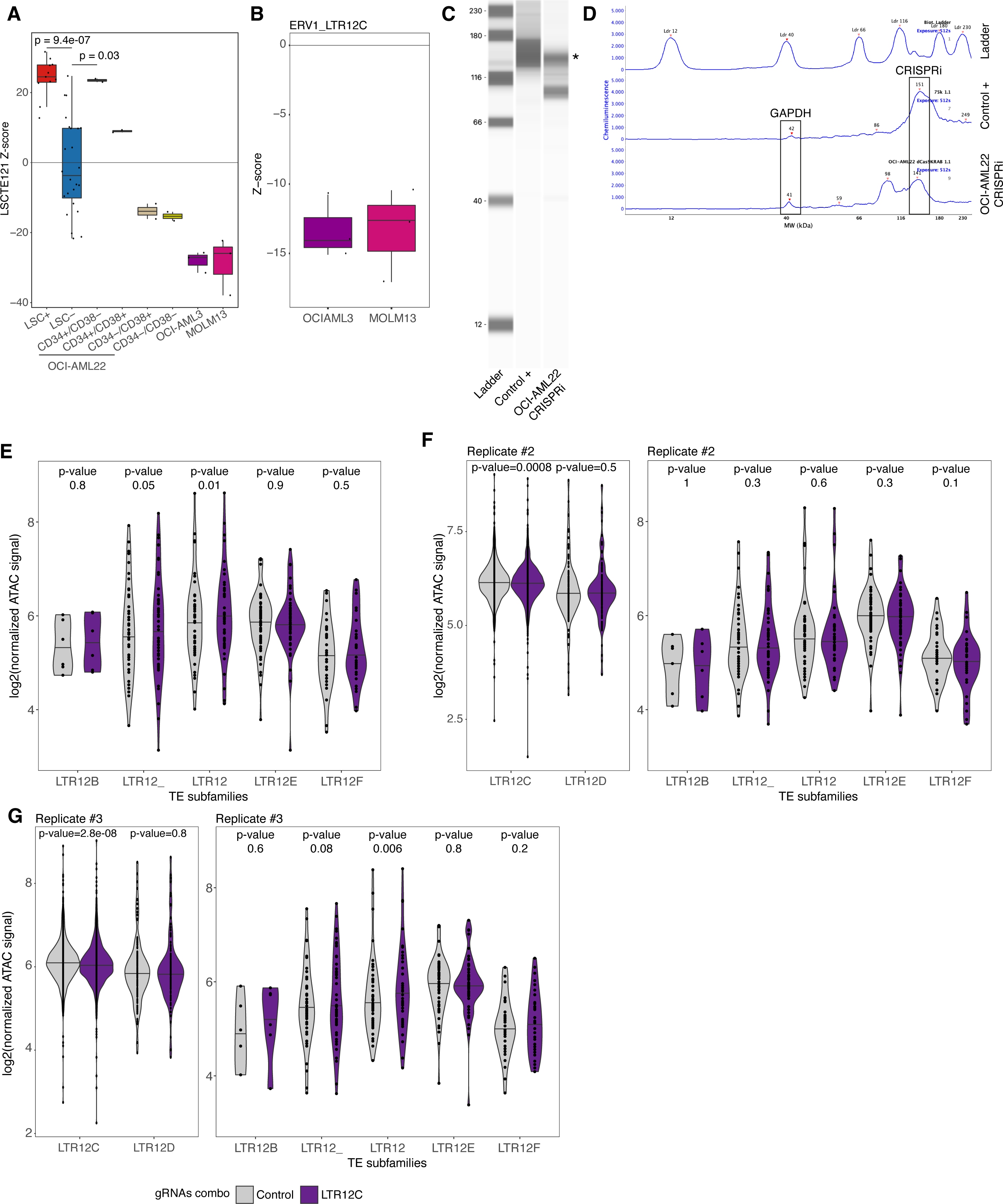
A) Box plot displaying the LSCTE121 Z-score in LSCs, OCI-AML22 fractions, OCI-AML3 and MOLM13 cell lines. p-values results of the Wilcoxon test are showcased on the box plots. B) Box plot displaying the LTR12C accessibility Z-scores in OCI-AML3 and MOLM13 cell lines. C) dCas9-KRAB expression levels in OCI-AML22 cells. Protein Simple (western blot) was performed on whole extracts, showcasing dCas9-KRAB expression in positive control ^29^. The star indicates the dCas9-KRAB band in CRISPRi OCI-AML22. D) Quantification of the chemiluminescence signal presented in C. Signal peaks corresponding to GAPDH (loading control) and dCas9-KRAB (CRISPRi) are indicated in the Fig.. E) Violin plots showcasing the chromatin accessibility signal distribution in replicate #1 over all LTR12 TE subfamilies, except LTR12C and LTR12D (in Fig. 5E). Every dot represents an individual TE element. p-values results of paired samples Wilcoxon test are showcased on the violin plot. F) Violin plots showcasing the chromatin accessibility signal distribution in replicate #2 OCI-AML22-LSC+ stably expressing both CRISPRi and scramble (control in gray) or LTR12C (purple) guide RNA combos, over LTR12C or LTR12D (left) and all other LTR12 TE subfamily elements (right). Every dot represents an individual TE element. Note that the loss of chromatin accessibility is specific to LTR12C elements only. p-values results of paired samples Wilcoxon test are showcased on the violin plot. G) Same as D but for replicate #3 OCI-AML22-LSC+ stably expressing both CRISPRi and scramble or LTR12C guide RNA combos.

